# Assessment of risk to hoary squash bees (*Peponapis pruinosa*) and other ground-nesting bees from systemic insecticides in agricultural soil

**DOI:** 10.1101/434498

**Authors:** D. Susan Willis Chan, Ryan S. Prosser, Jose L. Rodríguez-Gil, Nigel E. Raine

**Affiliations:** School of Environmental Sciences, University of Guelph, Guelph, Ontario, N1G 2W1, Canada; Department of Biology, University of Ottawa, Ottawa, Ontario, K1N 6N5, Canada

**Keywords:** crop pollination, environmental exposure distribution, insect pollinators, neonicotinoid insecticide, probabilistic risk assessment, solitary bees, systemic pesticides

## Abstract

Using the hoary squash bee (*Peponapis pruinosa*) as a model, we provide the first probabilistic risk assessment of exposure to systemic insecticides in soil for ground-nesting bees. To assess risk in acute and chronic exposure scenarios in *Cucurbita* and field crops, concentrations of clothianidin, thiamethoxam and imidacloprid (neonicotinoids) and chlorantraniliprole (anthranilic diamide) in cropped soil were plotted to produce an environmental exposure distribution for each insecticide. The probability of exceedance of several exposure endpoints (LC_50_s) was compared to an acceptable risk threshold (5%). In *Cucurbita* crops, under acute exposure, risk to hoary squash bees was below 5% for honey bee LC_50_s for all residues evaluated but exceeded 5% for clothianidin and imidacloprid using a solitary bee LC_50_. For *Cucurbita* crops in the chronic exposure scenario, exposure risks for clothianidin and imidacloprid exceeded 5% for all endpoints, and exposure risk for chlorantraniliprole was below 5% for all endpoints. In field crops, risk to ground-nesting bees was high from clothianidin in all exposure scenarios and high for thiamethoxam and imidacloprid under chronic exposure scenarios. Risk assessments for ground-nesting bees should include exposure impacts from soil and could use the hoary squash bee as an ecotoxicology model.

## Introduction

Global insect pollinator declines are being driven by multiple interacting environmental stressors, including land-use intensification, pathogens, invasive species and climate change, and may threaten the production of crops that depend directly or indirectly on the pollination services that bees provide^1,2^. For bee populations living in proximity to agricultural production, exposure to pesticides is one of the major environmental stressors likely affecting population health^2,3^.

*Cucurbita* crops (e.g., pumpkin, squash, summer squash, and gourds) are grown globally for their fruits. Because of their imperfect flowers and heavy, oily pollen, they are dependent upon bees to mediate pollination^4^. The insecticides used to control pests in *Cucurbita* crops (including the cucumber beetle, *Acalymma vittatum*) can also harm beneficial insect pollinators, setting up a tension between the need to control pests while maintaining the health of bee populations for the essential pollination services they provide.

In Ontario, three neonicotinoids (imidacloprid, clothianidin, thiamethoxam) are commonly used in *Cucurbita*-crop production to control insect pests^5^. Although they are effective against pests, neonicotinoid insecticides are of environmental concern because of their relatively high toxicity to (non-target) insects, their systemic nature, their persistence, and their extensive use in agriculture^6,7^. In agriculture globally, about 60% of neonicotinoids are applied as seed coatings or as in-furrow soil applications, with the remaining applied as foliar sprays^6^. Neonicotinoid residues have been found in the nectar and pollen of *Cucurbita* flowers^8^.^9^, and in agricultural soil^3,7^, where they have been found to persist in from season-to-season^7,10,11^.

For non-*Apis* bees, all developmental stages (adult and larvae) may be exposed to pesticide residues in consumed nectar and pollen. The extent of exposure is likely greatest for adult females because they consume pollen and nectar during sexual maturation and egg laying^12-14^, consume nectar to fuel their foraging and nesting activities, and handle pollen and nectar to feed their offspring^13,15^. Males consume nectar and pollen during sexual maturation, and nectar thereafter to fuel flight^12^. Larvae consume and topically contact pollen and nectar in their larval provisions^12^. For non-*Apis* bees, exposure may also be via nesting materials^14,16^. For ground-nesting bees, exposure from nesting sites is via soil contacted during nest excavation and construction for adult females, and via contact with the soil that forms nest cells during larval development. However, exposure within nest cells may be precluded because of the water-resistant coating applied to nest cells by many ground-nesting species^12^. Adult male ground-nesting bees have little exposure to soil as they do not participate in nest construction^12^. The persistence of neonicotinoids in soil makes both acute and chronic contact exposure scenarios for ground-nesting bees plausible. Although neonicotinoid uptake from soil by bees has not yet been quantified, the translocation of neonicotinoids to bees from residues in dust generated during corn planting is well documented and may be an appropriate parallel^17^.

Although their impacts on most wild bees are unknown, impacts of exposure to neonicotinoids on managed social and solitary bees include sublethal effects at sub-cellular to population levels, and at both adult and larval stages^18-20^. Substantial knowledge gaps remain around the toxicity and effects of neonicotinoids to arthropods in soil, including ground-nesting bees. Furthermore, exposure routes relating to nesting materials, including soil for ground-nesting bees, are not considered as part of current, honey bee centric, regulatory risk assessments for pesticide impacts on pollinators^14^.

Hoary squash bees (*Peponapis pruinosa*) are solitary bees that build their nests in the ground (Fig. 1) within *Cucurbita* cropping areas^21^ and consume mostly *Cucurbita*-crop pollen and nectar^22^. In eastern North America, hoary squash bees are one of the most important pollinators of *Cucurbita* crops^23,24^ and are obligately associated with these crops because they lack a wild plant host^25^. In 2014, the Pesticide Management Regulatory Agency (PMRA) of Health Canada initiated a special review of registered neonicotinoid insecticides used for pest control on cucurbit crops because of concerns about their potential impacts on hoary squash bees^26^. The hoary squash bee is common, nests in (sometimes large) aggregations, is a dietary specialist, has a well-documented natural history^15,22,23,27^, and is easy to maintain in captivity (DSWC personal observation), making it a promising candidate for a model species to evaluate risk of exposure to pesticides in soils for other ground-nesting solitary bee species. This study is the first to evaluate risk of exposure to insecticides in soil for ground-nesting bees. Our aims are (1) to evaluate which insecticides and exposure matrices (soil, pollen, nectar) pose a potential hazard to hoary squash bees; (2) to determine which hoary squash bee developmental stage (adult female or larvae) is at greatest hazard; (3) to evaluate risk from agricultural soil for ground-nesting solitary bees; and (4) to contribute to risk assessments for hoary squash bees and other solitary bees.

**Figure 1.**
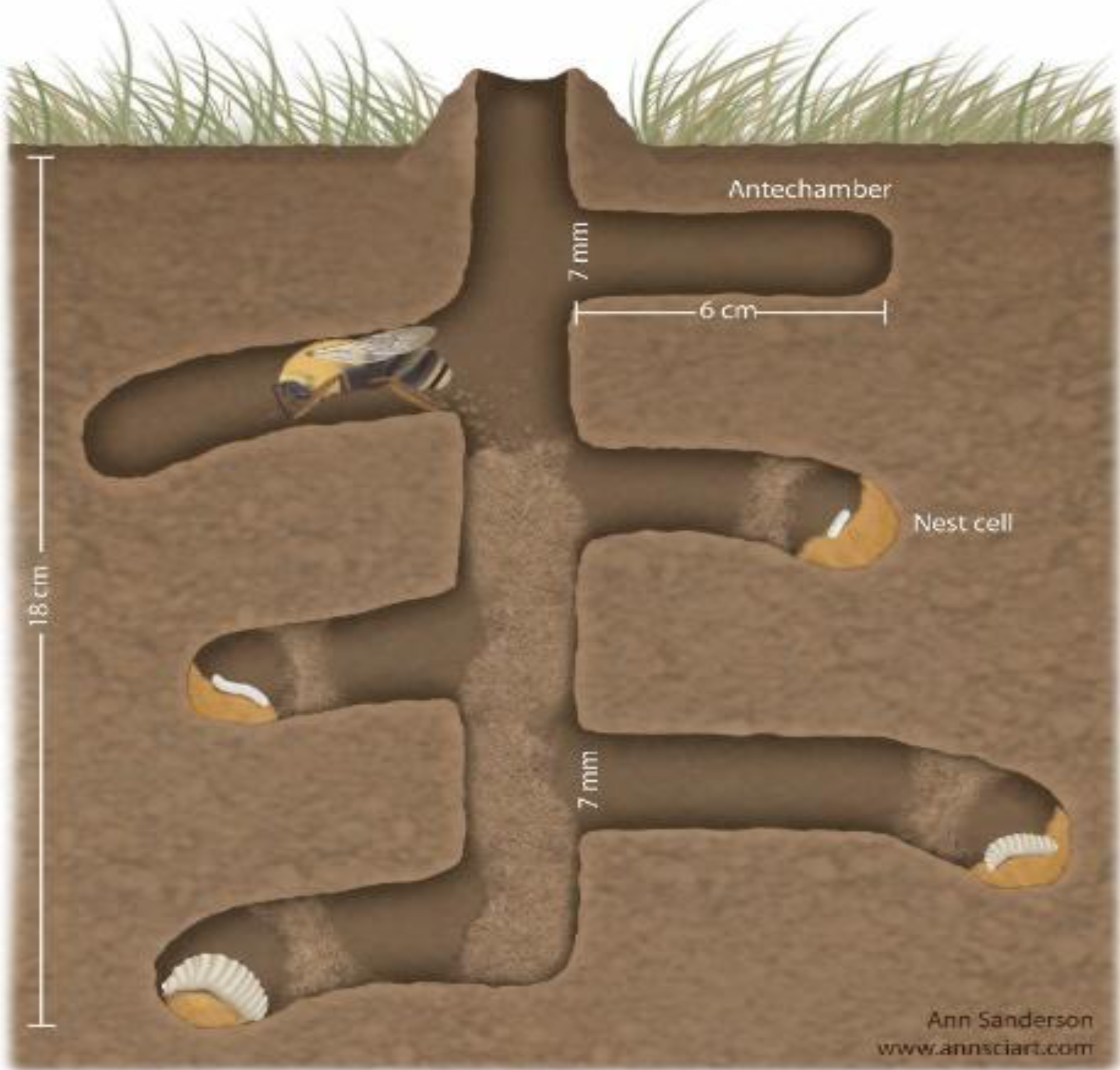
Nest of a hoary squash bee (*Peponapis pruinosa*) showing an adult female excavating a lateral tunnel and 4 immature stages (larvae) in sealed nest cells. Each nest cell is coated with a water-resistant lining. Soil from the main tunnel is moved to the soil surface and soil from lateral tunnels is backfilled into the vertical tunnel. The length of lateral tunnels varies. Graphic produced by Ann Sanderson and owned by authors DSWC and NER.

## Methods

In 2016, 29 samples of soil, nectar and pollen were taken from 18 *Cucurbita*-crop fields across southern Ontario. Ten soil samples (15 cm deep) were taken per field, combined and subsampled to produce a single 3 g sample for residue analysis. Twenty-five pollen and nectar samples were collected directly from staminate flowers. Nectar was harvested into a single 2 mL microcentrifuge tube using a 20 µL micro-pipette. Pollen was scraped off anthers from the same flowers, weighed, and put into a 2 mL tube. To determine the number of pollen grains per staminate flower, forty full anthers were gathered individually into 2 mL microcentrifuge tubes to which 0.5 mL of 70% alcohol was added. Pollen was dislodged from anthers by centrifuging at 2500 rpm for 3 minutes, the anther was removed, and the tubes were topped up to 2 mL with 50% glycerin solution. The suspension was mixed with a mini vortex mixer and the number of pollen grains in five 5 µL aliquots were counted on a grid under 25x magnification. The mean number of pollen grains per 5 µL aliquot was then related back to the full 2 mL volume. The mass of a single *Cucurbita* pollen grain was determined by dividing the mean mass of pollen grains per anther (mean ± sd = 0.0302 ± 0.0211 g; n = 25) by the mean number of pollen grains per anther (mean ± sd = 18438 ± 9810; n = 40) (0.0302 g / 18438 pollen grains = 1.64 × 10^−6^ g/pollen grain).

Information from the literature and this study were used to determine the realistic amount of pollen, nectar, and soil that female hoary squash bees would be exposed to via contact or ingestion or larvae would be exposed to via ingestion (Table S2). Males were not included in the evaluation, but are likely less exposed to pollen, nectar, and soil than females because they do not provision or construct nests^15^. The amount of pollen consumed by larvae (0.0542 g pollen/nest cell) was calculated by multiplying the mass of a single pollen grain (as calculated above = 1.64 × 10^-6^ g/pollen grain) by the mean number of pollen grains in hoary squash bee larval provisions (33045.4 ± 12675.2 pollen grains)^27^. Larval exposure to pesticide residues via contact with pollen in provisions was not evaluated. Contact exposure for adult females was assessed to be five times that for each larva because on average each female provisions five cells within a nest^15^. The amount of pollen and nectar ingested by adult females is unknown.

However, for pollen-collecting honey bee workers, which are most like adult female hoary squash bees in their foraging behaviour, nectar consumption has been estimated at 10.4 mg sugar/day to fly between their nest and forage patches and from flower-to-flower within those patches^40^. Based on this and an assumption of 40% sugar by volume in *Cucurbita pepo* nectar^27^, female hoary squash bees would consume 312 mg sugar in 780 mg of *Cucurbita* nectar to meet their energy requirements over 30 days. This is likely an over estimation as the foraging radius of squash bees from nest to flower patch is much smaller than that of honey bees (honey bee foraging radius: mean = 5.5, maximum = 15 km^58^; oligolectic solitary bee foraging radius <260 m^54^). Using hoary squash bee nest dimensions from Mathewson^15^, the volume of soil excavated by a female squash bee to build a nest is 25.19 cm^3^ (Table S1). Multiplying the volume by the bulk density (BD) of loam soil, a common agricultural soil (BD_loam_ = 1.33 g/cm^3^)^59^, gives the mass of soil that a female hoary squash bee contacts as she constructs a nest with 5 nest cells over 30 days (25.19 cm^3^ x 1.33 g/cm^3^ = 33.51 g). Acute (48 h) exposure was calculated by dividing cumulative exposure by 15 (33.5g / 15 = 2.23 g). Table S2 summarizes exposure routes and extent of exposure for the hoary squash bee.

All samples collected from *Cucurbita* farms were submitted for analysis to University of Guelph Agri-Food Laboratories (Table S3) (ISO/IEC 17025 accredited). Samples were analyzed using their TOPS-142 LC pesticide screen, modified from the Canadian Food Inspection Agency (CFIA) PMR-006-V1.0 method. Pesticides were extracted using the QuEChERS Method^60^.

Extracts were analyzed using high performance liquid chromatography paired with electrospray ionization and tandem mass spectrometry and gas chromatography paired with tandem mass spectrometry.

Honey bee lethal effect endpoints (e.g., the concentration causing 50% mortality, LD_50_ or LC_50_) were used for hoary squash bees because current regulatory standards consider the honey bee to be an adequate proxy for all bee species^14,31^ and because lethal doses for larval stages are rarely available^18,20^. Honey bee LD_50_ and LC_50_ values were obtained from the US-EPA Pesticide Ecotoxicity Database of the Office of Pesticide Programs, Ecological Fate and Effects Division, of the U.S. Environmental Protection Agency^61^ and published literature. A geometric mean of multiple LC_50_ values, reported from many sources^35,61-66^ and the lowest reported LC_50_ value were used in this study. For adult female hoary squash bees contacting soil and pollen during nest construction and provisioning, contact honey bee LC_50_ values were used. Oral honey bee LD_50_ values were used for adult female hoary squash bee ingestion of nectar, and larval hoary squash bee ingestion of pollen. For probabilistic risk assessment of soil exposure, the same lethal effect endpoints and a surrogate solitary bee contact LC_50_ (honeybee LC_50_/10)^32,67^ were used, and lethal effect endpoints were converted to soil exposure concentrations as follows:

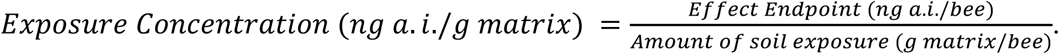

Hazard quotients (HQ) were calculated for each residue detected in each exposure matrix as follows:

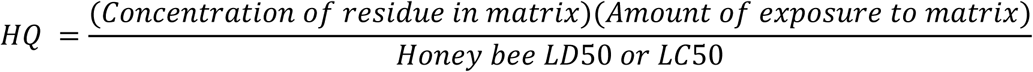

An HQ >1 indicates that a pesticide is a potential lethal hazard. For HQs, the residue concentrations were calculated by taking the geometric mean of all quantifiable concentrations in samples. Concentrations of residue were reported in ng a.i./g matrix, amount of exposure to a matrix was reported in g matrix/bee, and LD_50_s or LC_50_s were reported in ng a.i./bee. Hazard quotients were summed for each type of pesticide (fungicide or insecticide), each exposure matrix (soil, pollen, nectar), and each developmental stage (adult female, larvae). After determining HQs, the hoary squash bee exposure models for both acute (48 h, 2.23 g soil) and chronic (30 days, 33.5 g soil) scenarios were used to carry out probabilistic risk assessment for both the hoary squash bee in *Cucurbita* crop soils using data from our own samples and for ground-nesting solitary bee species generally in field crops soils using a publicly-available dataset provided by the Ontario government^11^. The government data set reported neonicotinoid residues in soil (15 cm depth) from 38 agricultural sites in southern Ontario in 2016 with a limit of detection of 0.05 ng/g for clothianidin, imidacloprid, and thiamethoxam. Insecticide residues in our samples represented field-realistic exposure for hoary squash bees on *Cucurbita* farms in 2016. Neonicotinoid residues in the MOECC samples represented persistent residues in soil from a previous cropping cycle, rather than active ingredients applied during the 2016 season. Environmental exposure distributions (EEDs) were constructed for each neonicotinoid in each exposure scenario and each crop type. Chronic exposure scenarios are reasonable for neonicotinoids because their effect on the nicotinic acetylcholine receptors (nAChRs) of insects are cumulative and irreversible^68^. Although an EED for chronic exposure to chlorantraniliprole was constructed, evidence from honey bees suggests its effects may be transient as honey bees dosed orally or topically under artificial test conditions became lethargic but recovered within 48-72 hours^69^.

EEDs were generated from the concentration of insecticides measured in soil by fitting the data to a log-normal or gamma distribution via maximum likelihood estimation (MLE), using the fitdistrcens function in the R package fitdistrplus^70-71^. This function allows for the fit of censored (right-, left-, or interval-censored) data enabling the use of the available “non-detect” (samples containing residues below the limit of detection (LOD)), and interval (samples containing residues above the limit of detection (LOD) but below the limit of quantification (LOQ)) data. The fitdistrplus function calculates the probability plotting position using Hazen’s rule, with probability points of the empirical distribution calculated as (1:n - 0.5)/n, where n is the total number of data points^70^. Interval data (non-detects, LOD-LOQ) were ranked according to the midpoint of the interval. Confidence intervals (95%) for the distribution parameters and distribution estimates were calculated via nonparametric bootstrapping (1000 iterations) with the bootdiscens function in the fitdistrplus package^70^. The fit to other distributions (Weibull and Exponential) were also tested, and the best fit was chosen via comparison of the Akaike information criterion (AIC) supported by visual inspection of the fit to the actual data. For data from *Cucurbita* crop soils, the gamma distribution provided the best fit to construct EEDs for clothianidin and chlorantraniliprole, and the log-normal distribution provided the best fit for imidacloprid. EEDs were not constructed for thiamethoxam as there was no quantifiable residue data. For data from field crops soils, the gamma distribution provided the best fit for all residues. Table S7 provides model parameters for all EEDs. The percent rank was subtracted from 100 to determine the percent exceedance which is the probability that a population will be exposed to a benchmark lethal dose. The benchmarks used were exposure concentrations associated with the lethal effect endpoints described (geomean LC_50_, lowest LC_50_, solitary bee surrogate LC_50_: Table S5). Table S6 presents exceedances at 100% translocation of residues from the soil to the bee, however recognizing that translocation is likely lower than 100%, exceedances at 10, 25, 50, 75% translocation are also presented in Table S8. Exceedance lower than 5% was considered acceptable risk because it assures 95% protection of the population.

## Results

### Pesticide Residue Profiles-*Cucurbita* Crops

Residues of 7 insecticides, 6 fungicides, and 2 herbicides were detected in samples taken from *Cucurbita* crops: pesticides detected, limits of detection and quantification (LOD/LOQ), frequency of detection, and mean and maximum concentrations in each exposure matrix are shown in Table S3 (herbicide residues were not assessed further in this study). The three exposure matrices (pollen, nectar, soil) show different pesticide residue profiles (Table S3). In soil, 5/7 insecticides and 5/6 fungicides were detected. Pollen contained residues of 4/7 insecticides and 6/6 fungicides, but residues were detected much less frequently and at lower concentrations than in soil. Nectar contained 3/7 insecticides and 3/6 fungicides, with the lowest residue concentrations and frequency of detection. Imidacloprid was the only insecticide detected in all three matrices (in 21% of soil samples and 3% of pollen and nectar samples).

Thiamethoxam was present only in a single soil sample, at a concentration below the limit of quantification. Clothianidin was detected in 34% of soil samples but was not detected in nectar or pollen. The insecticides chlorantraniliprole and carbaryl were detected in soil (in 24% and 10% of samples respectively) and pollen samples (3% and 7% respectively), but not in nectar. The insecticides methomyl and dimethoate were not detected in soil, but dimethoate was detected in pollen and nectar (3% each), and methomyl was detected in 7% of nectar samples. The fungicides pyraclostrobin and propamocarb were detected in all three matrices, whereas boscalid, quinoxyfen and difenoconazole were detected in soil and pollen only, and picoxystrobin was detected in nectar and pollen only.

### Hazard Assessment-Hoary Squash Bee

#### Pesticides

Hazard quotients (HQ) for fungicide and insecticide residues in soil, pollen, and nectar for adult females and larvae are presented in Table 1. As expected, the combined HQ for insecticides (HQ_insecticide total_ = 4.92) was much higher in all exposure matrices than fungicides (HQ_fungicide_ = 0.03). Among the insecticides, only clothianidin and imidacloprid had HQs ≥ 1 in soil, and the combined HQ of all non-neonicotinoid insecticides in soil was low (HQ_non-neonicotinoid_ = 0.06).

**Table 1.** Hazard quotients (HQ) for each active ingredient, and pesticide type (insecticide, fungicide) in three exposure matrices (soil, pollen, nectar) and both developmental stages (adult female or larvae) based on honey bee LD_50_ values (taken from US-EPA database, unless otherwise noted), and mean residue concentration in the exposure matrices for all pesticide residues found detected on 18 *Cucurbita-*crop farms in Ontario in 2016. Numbers shown in bold face are combined hazard quotients for a pesticide type, an exposure matrix, or a developmental stage. Where “ND” is indicated, residues were not detected, where “NQ” is indicated, residues were not quantifiable in samples. a.i. = active ingredient.

|  | LC<sub>50</sub> Endpoint<br>(Honey bee) |  | SOIL |  | POLLEN |  |  | NECTAR |  | COMBINED HAZARD<br>QUOTIENT |  |  |
| --- | --- | --- | --- | --- | --- | --- | --- | --- | --- | --- | --- | --- |
|  | Contact | Oral | Mean<br>Conc.<br>in Matrix | HQ<br>Adult<br>female | Mean<br>Conc.<br>in Matrix | HQ Larvae<br>(Oral) | HQ<br>Adult Female<br>(Contact) | Mean<br>Conc. in<br>Matrix | HQ<br>Adult Female | Across<br>Exposure<br>Matrix,<br>All Stages | Adult<br>Female | Larvae |
|  | ng<br>a.i./bee | ng a.i./bee<br>or ppb | ng a.i./g | =33.5 g/bee<br>*Matrix<br>Conc. /LD<sub>50</sub> | ng a.i./g | =0.0542 g/bee<br>* Matrix<br>Conc. /LD<sub>50</sub> | =5*0.0542<br>g/bee* Matrix<br>Conc. /LD<sub>50</sub> | ng a.i./g | =0.78 g/bee*<br>Matrix Conc.<br>/LD<sub>50</sub> | ΣHQ | ΣHQ | ΣHQ |
| INSECTICIDE ΣHQ |  |  |  | 4.38 |  | 0.06 | 0.03 |  | 0.45 | 4.92 | 4.86 | 0.06 |
| Neonicotinoid ΣHQ |  |  |  | 4.32 |  | 0.06 | 0.03 |  | 0.18 | 4.59 | 4.53 | 0.06 |
| Clothianidin (geomean) <sup>A</sup> | 35.88 | - | 1.95 | 1.82 | ND | - | 0 | ND | 0 | 1.82 | 1.82 | 0 |
| Imidacloprid (geomean) <sup>B</sup> | 40.03 | 3.9 | 2.99 | 2.50 | 4.3 | 0.06 | 0.03 | 0.88 | 0.18 | 2.76 | 2.71 | 0.06 |
| Thiamethoxam (geomean) <sup>C</sup> | 25.64 | - | NQ | 0 | ND | 0 | 0 | ND | 0 | 0 | 0 | 0 |
| Chlorantraniliprole <sup>D</sup> | >81500 | >117800 | 36.82 | 0.02 | 68 | 1.69E-5 | 2.26E-4 | ND | 0 | 0.02 | 0.02 | 2.26E-4 |
| Carbaryl <sup>E</sup> | 11200 | - | 14.2 | 0.04 | 16.47 | - | 03.9E-4 | ND | 0 | 0.04 | 0.04 | 3.99E-4 |
| Dimethoate <sup>E</sup> | - | 56 | ND | 0 | 6.2 | 0 | 0 | 0.5 | 0.01 | 0.01 | 0.01 | 0 |
| Methomyl (ppb) <sup>E</sup> | - | 1.18 | ND | 0 | ND | 0 | 0 | 0.39 | 0.26 | 0.26 | 0.26 | 0 |
| FUNGICIDE ΣHQ |  |  |  | 0.03 |  | 1.24E-4 | 6.18E-4 |  | 9.20E-5 | 0.03 | 3.08E-2 | 1.24E-4 |
| Pyraclostrobin <sup>E</sup> | 100000 | - | 3.8 | 1.27E-4 | 29.65 | - | 8.04E-5 | 2 | - | 1.35E-3 | 1.35E-3 | - |
| Picoxystrobin <sup>E</sup> | 200000 | - | ND | 0 | 4.55 | - | 6.17E-6 | 0.3 | - | 6.17E-6 | 6.17E-6 | - |
| Boscalid <sup>E</sup> | 200000 | 166000 | 46.22 | 7.74E-3 | 17.82 | 5.28E-6 | 2.41E-5 | ND | 0 | 7.77E-3 | 7.77E-3 | 5.28E-6 |
| Propamocarb <sup>E</sup> | 100000 | 116000 | 23.03 | 7.72E-3 | 222.06 | 1.04E-3 | 6.02E-4 | 11.18 | 7.52E-5 | 9.43E-3 | 8.39E-3 | 1.04E-3 |
| Quinoxifen <sup>E</sup> | 100000 | 1000000 | 7.86 | 2.63E-3 | 79.14 | 4.29E-5 | 2.14E-4 | ND | 0 | 2.85E-3 | 2.85E-3 | 4.29E-5 |
| Difenoconazole <sup>E</sup> | 100000 | 177000 | 18.87 | 6.32E-3 | 16.46 | 5.04E-5 | 4.46E-5 | ND | 0 | 6.37E-3 | 6.37E-3 | 5.04E-5 |

<sup>A</sup>35,60,62
<sup>B</sup>35,60,62,63,64
<sup>C</sup>60,62,65
<sup>D</sup>68
<sup>E</sup>60

Although chlorantraniliprole had an HQ <1, it was included in further probabilistic risk assessment for comparison because, like the neonicotinoids, it is systemic and was present in 24% of samples. The physical, chemical and environmental fate properties of imidacloprid, clothianidin, thiamethoxam and chlorantraniliprole are presented in Table S4.

#### Exposure Matrices

Hazard quotients for pollen, including both adult and larval exposure (HQ = 0.36), and nectar (HQ = 0.45) were low relative to HQs for soil (Table 1). However, only adult female exposure was assessed for nectar as little information exists about the amount of nectar consumed by larvae or adult male solitary bees. Low HQs for pollen and nectar reflect the low concentrations of all the insecticides detected in these exposure matrices. Imidacloprid was the largest contributor to hazard for pollen, and methomyl was the largest contributor for nectar (Table 1). As HQ <1 for pollen and nectar, they were deemed non-hazardous for the lethal dose endpoint. This does not imply that there are no hazards associated with sublethal endpoints as these were not evaluated.

The combined HQ for insecticides in soil was high (HQ_soil_ = 4.38: Table 1). Soil HQ was mostly attributable to the neonicotinoid residues (HQ_combined soil neonicotinoids_ = 4.32; 4.32/4.38=99%).

#### Developmental Stage

Hazard is greater for adult females (HQ_adult female_ = 4.86) than for larval hoary squash bees (HQ_larvae_ = 0.06), mostly because of the adults’ exposure to neonicotinoid residues in soil during nest construction (HQ_soil,adult female_ = 4.32: Table 1).

### Probabilistic Risk Assessment

#### For Hoary Squash Bees in *Cucurbita* Crops

For the acute exposure scenario (2.23 g soil, 48 h), environmental exposure distributions (EEDs) indicate that clothianidin showed exceedance below 5% for the geometric mean honey bee LC_50_ and the lowest honey bee LC_50_ endpoints. However, exceedance for the solitary bee surrogate LC_50_ endpoint (28.3%) was relatively high (Fig. 2A). Imidacloprid showed exceedance below 5% for the geometric mean honey bee LC_50_ (3.5%) and exceedances greater than 5% for both the lowest honey bee LC_50_ (8.9%) and the solitary bee surrogate LC_50_ (31.2%) (Fig. 2B). For the chronic exposure scenario (33.5 g soil, 30 days), exceedance for clothianidin in soil was above 5% for all exposure endpoints in *Cucurbita* crops (geometric mean honey bee LC_50_ = 35.8%; lowest honey bee LC_50_ = 44.3%; solitary bee surrogate LC_50_ = 68.7%) (Fig. 3A). For imidacloprid, the exceedance for all lethal endpoints was also high under the chronic exposure scenario (geometric mean honey bee LC_50_ = 39.8%; lowest honey bee LC_50_ = 57.8%; solitary bee surrogate LC_50_ = 85.4%) (Fig. 3B). Exceedance under both the acute and chronic exposure scenarios for chlorantraniliprole was below 5% for all lethal dose effect endpoints assessed (Fig. 3C). Because high limits of detection and quantification (LOD/LOQ) rendered no quantifiable residue samples, EEDs could not be fitted to data for thiamethoxam.

**Figure 2.**
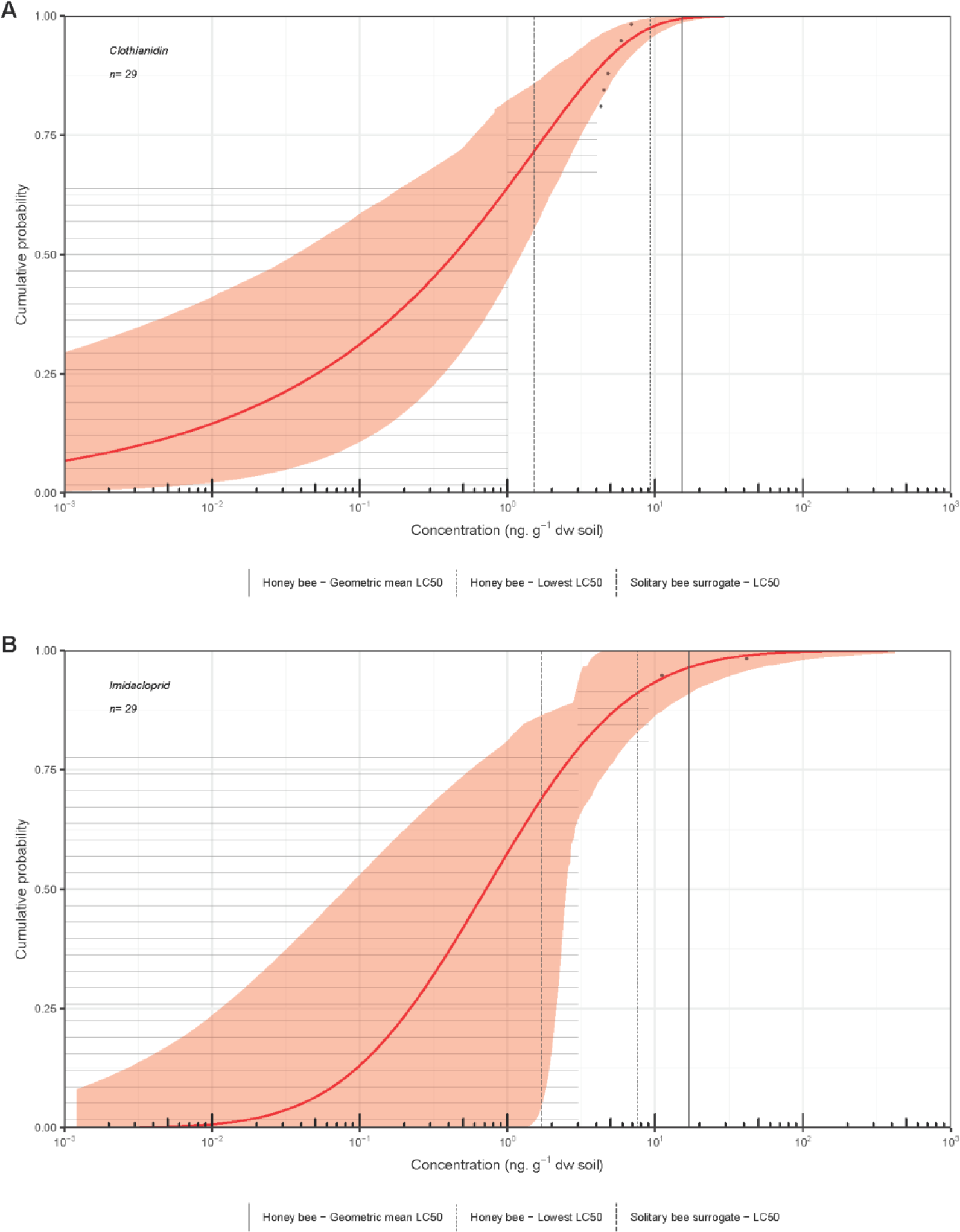
Environmental Exposure Distribution (EED) for acute exposure to (A) clothianidin and (B) imidacloprid concentrations measured in soil samples taken from 0-15 cm depth in *Cucurbita*-crop fields in Ontario, 2016. Effects benchmark concentrations for acute exposure (48 h, 2.23 g soil) for the hoary squash bee, *Peponapis pruinosa* (i.e. honey bee geometric mean LC_50_, honey bee lowest LC_50_, solitary bee surrogate LC_50_), are represented by vertical lines on the EED. Exceedance of these endpoints is calculated by subtracting the cumulative probability from one. Grey horizontal lines represent individual samples below the analytical limits of detection or quantification.

**Figure 3.**
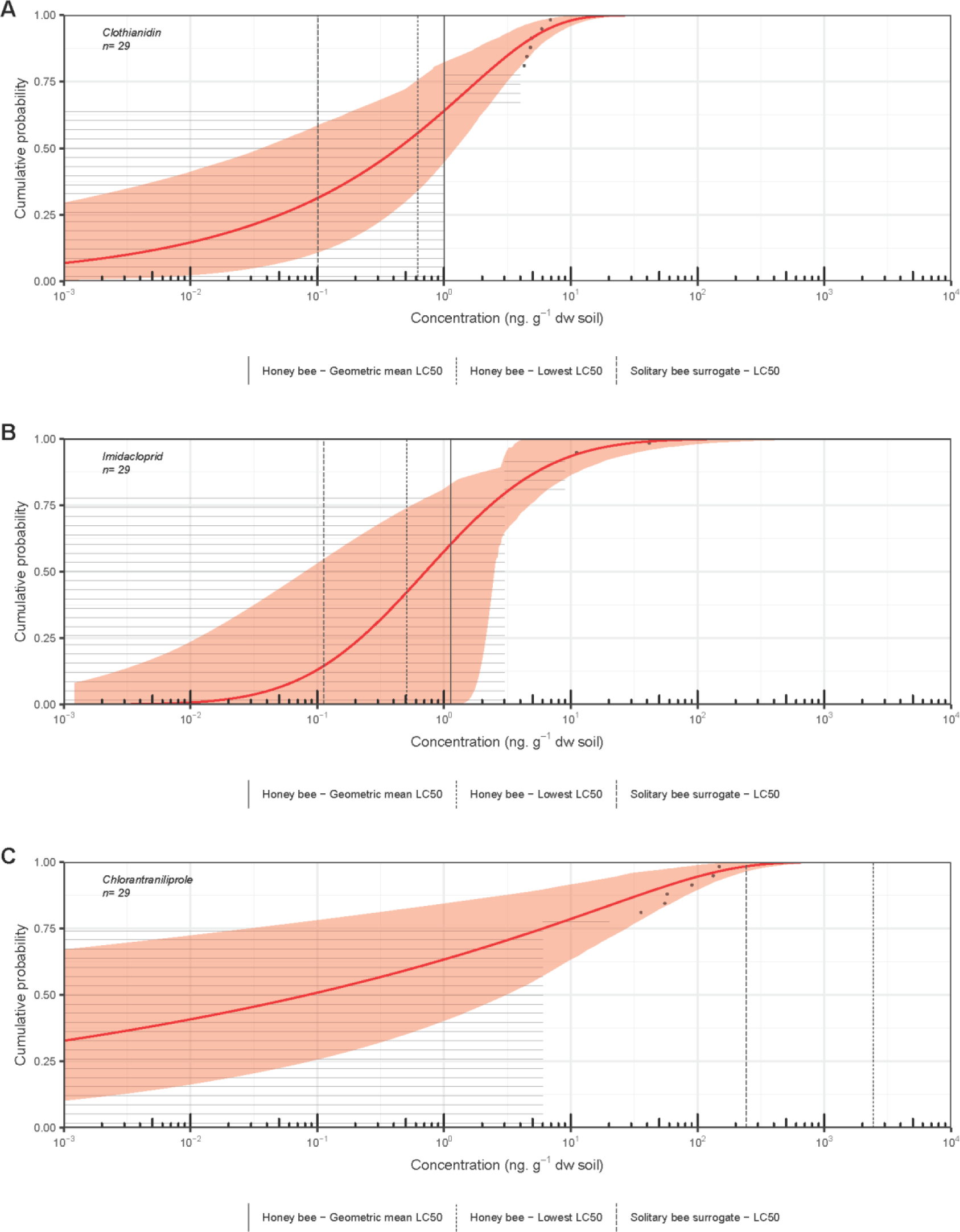
Environmental Exposure Distribution (EED) for chronic exposure to (A) clothianidin, (B) imidacloprid, and (C) chlorantraniliprole concentrations in soil samples taken from 0-15 cm depth in *Cucurbita*-crop fields in Ontario, 2016. Effects benchmark concentrations for chronic exposure (30 days, 33.5 g soil) for the hoary squash bee, *Peponapis pruinosa* (i.e. honey bee geometric mean LC_50_, solitary bee surrogate LC_50_), are represented by vertical lines on the EED. Exceedance of these endpoints is calculated by subtracting the cumulative probability from one. Grey horizontal lines represent individual samples below the analytical limits of detection or quantification.

#### For Ground-Nesting Bees in Field Crops

Clothianidin was detected in 96.34%, imidacloprid was detected in 10.97%, and thiamethoxam was detected in 81.48% of soil samples taken from Ontario field crops (n = 82)^11^. Hoary squash bee exposure amounts (2.23 g soil-acute exposure; 33.5 g soil-chronic exposure) were used as a surrogate for exposure to determine exceedance for ground-nesting bees generally. In the acute exposure scenario for imidacloprid (Fig. S1) and thiamethoxam (Fig. 4B), only the solitary bee surrogate LC_50_ endpoint showed exceedance above 5%. In contrast, for clothianidin the probability of exceedance was greater than 5% for all exposure endpoints (honey bee geometric mean LC_50_ = 11.72%; honey bee lowest LC_50_ = 27.25%; solitary bee surrogate LC_50_ = 81.85%), suggesting that risk to ground-nesting bees is high from clothianidin in field crops soils, even when exposure is acute (Fig. 4A). In the chronic scenario, probability of exceedance for clothianidin was very high for all exposure endpoints (honey bee geometric mean LC_50_ = 87.68%; lowest honey bee LC_50_ = 92.4%; solitary bee surrogate LC_50_ = 98.82%; Fig. 5A). For thiamethoxam, probability of exceedance was also high (honey bee geometric mean LC_50_ = 35.7%; honey bee lowest LC_50_ = 37.36%; solitary bee surrogate LC_50_ = 78.42%; Fig. 5B). Probability of exceedance for imidacloprid under the chronic exposure scenario was below 5% for the honey bee geometric mean LC_50_ (4.16%) and exceeded this threshold for the honey bee lowest LC_50_ (5.89%) and the solitary bee surrogate LC_50_ (9.24%) (Fig. S2; Table S6).

**Figure 4.**
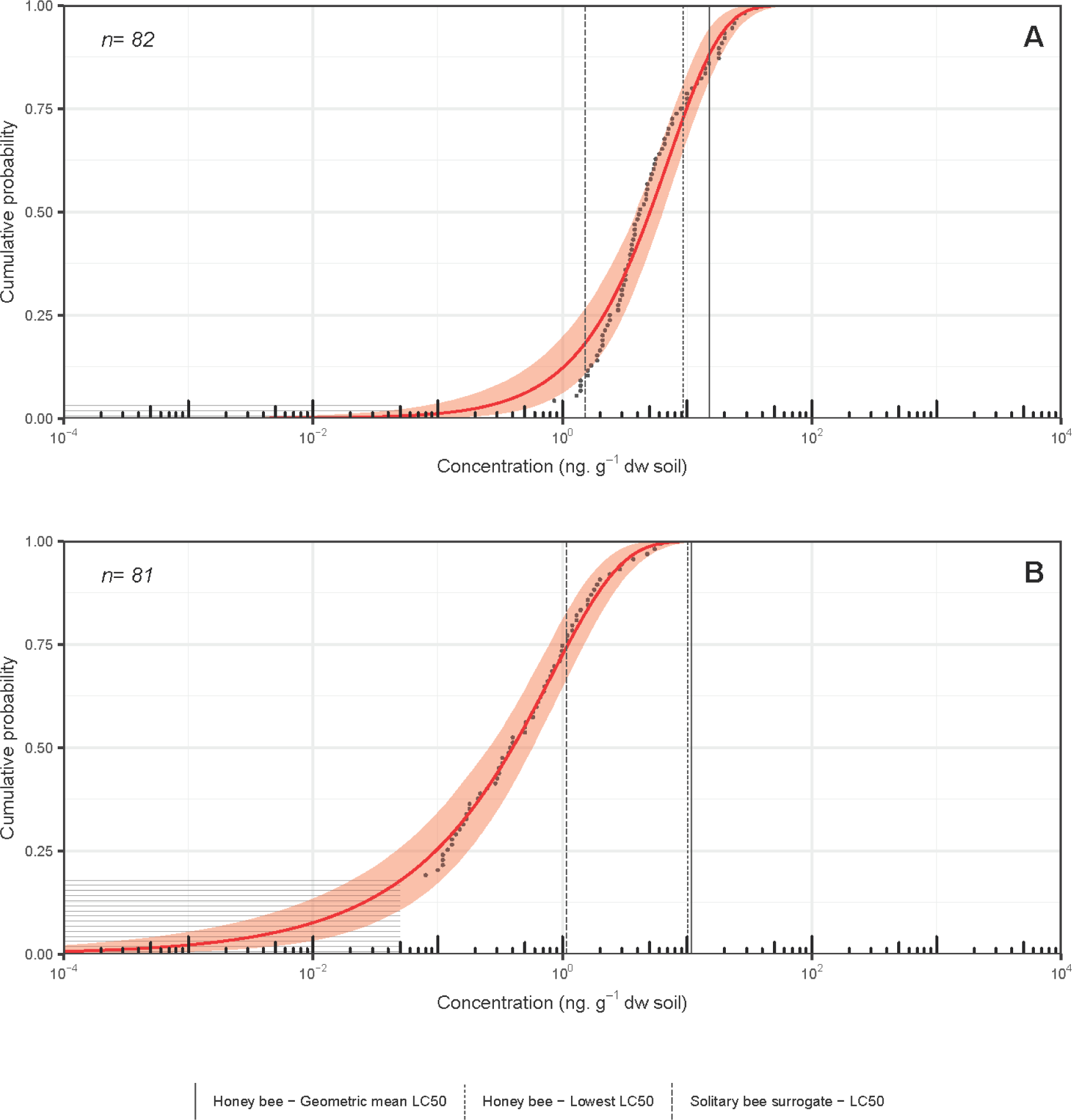
Environmental Exposure Distribution (EED) for acute exposure to (A) clothianidin and (B) thiamethoxam in soil from field crops (corn, soybeans, wheat) based on MOECC dataset^26^. Soil samples taken from 0-15 cm depth in southern Ontario, 2016. Effects benchmark concentrations are for solitary ground-nesting bees based on the hoary squash bee (*Peponapis pruinosa*) acute exposure amounts (48 h, 2.23 g soil). Effect benchmarks (i.e. honey bee geometric mean LC_50_, solitary bee surrogate LC_50_) are represented by vertical lines on the EED. Exceedance of these endpoints is calculated by subtracting the cumulative probability from one. Grey horizontal lines represent individual samples below the analytical limit of detection.

**Figure 5.**
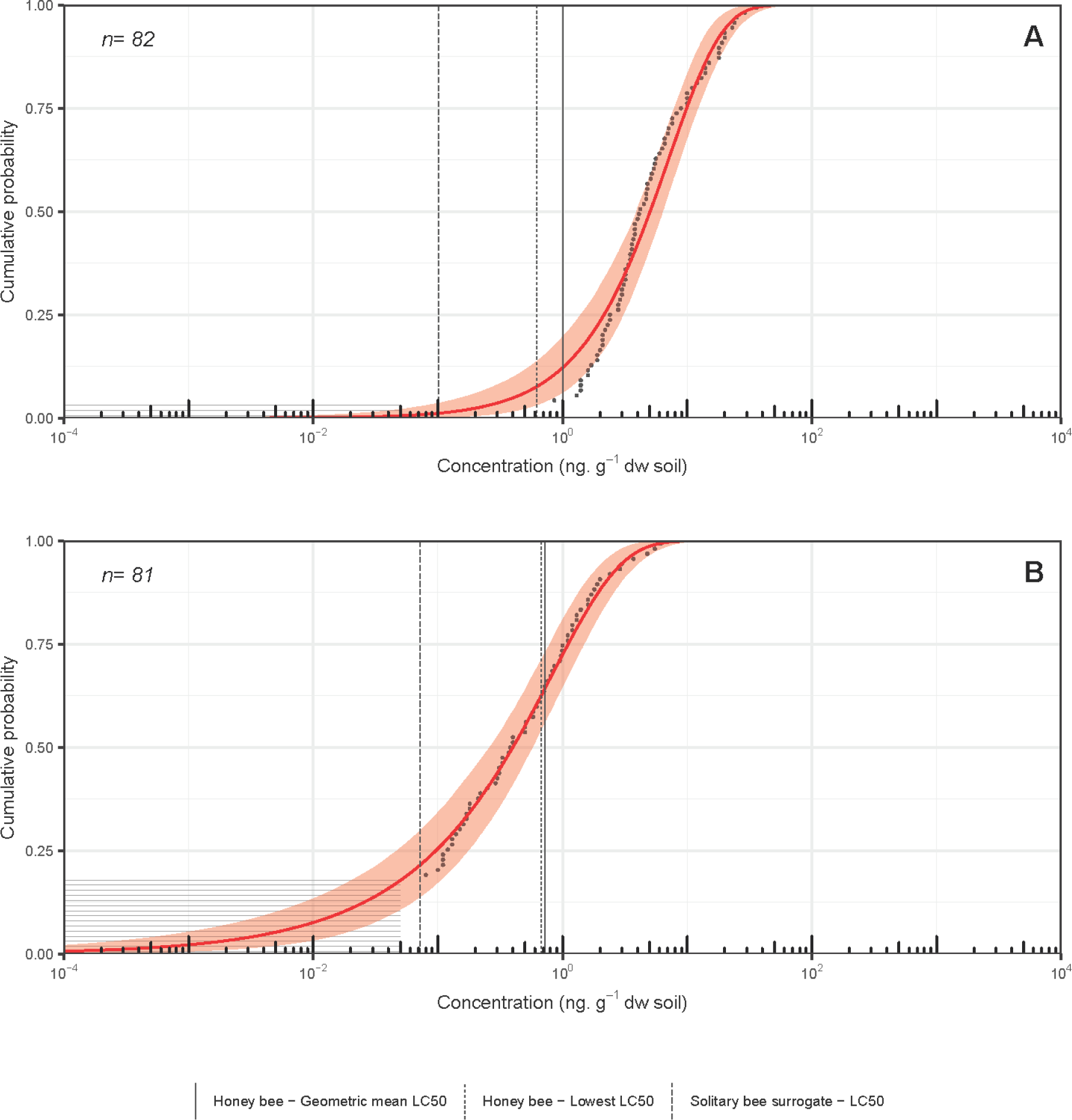
Environmental Exposure Distribution (EED) for chronic exposure (30 days, 33.5 g soil) to (A) clothianidin and (B) thiamethoxam in soil from field crops (corn, soybeans, wheat) based on MOECC dataset^26^. Soil samples taken from 0-15 cm depth in southern Ontario, 2016. Effects benchmark concentrations are for solitary ground-nesting bees based on the hoary squash bee (*Peponapis pruinosa*) exposure model. Effect benchmarks (i.e. honey bee geometric mean LC_50_, solitary bee surrogate LC_50_) are represented by vertical lines on the EED. Exceedance of these endpoints is calculated by subtracting the cumulative probability from one. Grey horizontal lines represent individual samples below the analytical limit of detection.

## Discussion

Comparing the three exposure matrices, soil had the greatest number of neonicotinoids detected (clothianidin, imidacloprid, thiamethoxam), whereas imidacloprid was the only neonicotinoid detected in pollen and nectar. The maximum concentration of imidacloprid detected in soil (41.6 ng/g) was substantially greater than in pollen (4.3 ng/g) or nectar (1.1 ng/g), because imidacloprid is applied directly to soil and can bind to soil particles^28^ (Table S3). Although thiamethoxam is more soluble than imidacloprid, it is quickly metabolized to clothianidin^29,30^, which is much less soluble than either imidacloprid or thiamethoxam (Table S4). Dively and Kamel^8^ reported both higher imidacloprid residue concentrations (60.9 ng/g for pollen, 7.4 ng/g for nectar) and greater frequencies of detection (92% of pollen samples; 88% of nectar samples) in the pollen and nectar of *Cucurbita* crops. However, the interval between application and sampling was shorter in their study (5 weeks) than in our study (8 weeks). Stoner and Eitzer^9^ also reported higher concentrations of imidacloprid and thiamethoxam in *Cucurbita* pollen and nectar than found in this study, likely because they included anther and nectary tissue in their samples. The analysis of the hazard of pesticides to the hoary squash bee within *Cucurbita* crops is based on LC_50_ for adult honey bees as this species is presently the regulatory standard for testing toxicity in bees^31^. It is possible that differences in sensitivity exist between these two species as not all bee species are equally sensitive to all pesticides^32-34^. Neonicotinoids are often more toxic to bees when exposure is via ingestion^18,19,35,36^, and solitary bees are more sensitive than either honey bees or bumble bees to oral exposure^33,34,36^, therefore the use of oral honey bee LD_50_ values in this study may under-represent oral toxicity to hoary squash bees. However, toxicity via contact exposure varies much less among species^32,37^, thus honey bee contact LC_50_ values may adequately represent contact toxicity for adult female hoary squash bees. The evaluation of hazard to larval squash bees using adult honey bee LD_50_ or LC_50_ values in this study may underestimate hazard because of species and developmental stage differences in sensitivity to neonicotinoids^18,19,29,38^.

Thiamethoxam appears to pose minimal hazard because residues were only detected in a single sample at concentrations below the limit of quantification. However, an absence of thiamethoxam residues may be the result of rapid metabolization to clothianidin, which was detected in our soil samples^29,30^. Both clothianidin (HQ = 1.82) and imidacloprid (HQ = 2.77) pose a hazard in *Cucurbita* growing systems in Ontario because they are detected frequently, and their respective HQs exceed one when summed across all exposure matrices. Imidacloprid appears to be more hazardous in this cropping system than clothianidin based on this deterministic approach because it was found at higher concentrations in soil and was also present in nectar and pollen (Table S3). The combined hazard of all neonicotinoids in the system was also high (HQ_combined neonicotinoid_ = 4.59), suggesting that hoary squash bee populations may be subject to hazards of exposure to lethal concentrations of neonicotinoids in a worst-case scenario. Combined hazard from chlorantraniliprole in all matrices was low (HQ <0.02), likely because it has a much higher LC_50_ than neonicotinoids.

There is common agreement that bees can be exposed to neonicotinoids from nectar and pollen in the flowers upon which they forage^16,18,19^. Although neither pollen nor nectar were deemed hazardous in this study based on their HQs, HQs were higher for adult hoary squash bees from insecticide exposure via *Cucurbita* nectar (HQ_nectar_ = 0.45) than via pollen (HQ_pollen_ = 0.03) because adult exposure to pollen was contact rather than oral and adult consumption of pollen was not evaluated, whereas adult exposure to nectar was oral and was likely an overestimation based on honey bee nectar consumption values. Contact lethal doses are higher than oral lethal doses for the residues detected (Table 1). Although no hazard from pollen or nectar was found for the lethal dose endpoint, sublethal effects are still possible at these low HQs.

Currently, there are no studies that evaluate the risks to ground-nesting bees from direct exposure to neonicotinoids in soil, although some studies assessing direct effects on other soil fauna exist^39^. Because both imidacloprid and thiamethoxam (which metabolizes to clothianidin) are applied to the soil in *Cucurbita* cropping systems, and may persist in the soil for longer than a single growing season in Canada^11^, it is unsurprising that the hazard to the ground-nesting hoary squash bee from neonicotinoids in soil (HQ_soil_ = 4.32) is much higher than even the combined hazard from neonicotinoids in both pollen and nectar (HQ_pollen+nectar_ = 0.27; Table 1). Therefore, soil appears to be the most important route of exposure to systemic pesticides for hoary squash bees.

The combined hazard from insecticides for adult female hoary squash bees from all exposure matrices (soil, pollen, nectar) was high, with 93% of this hazard attributable to neonicotinoids in soil (Table 1). Mathewson^15^ reported that hoary squash bees can construct more than one nest per season when environmental conditions (e.g. nectar and pollen resources, weather) permit.

However, in this study, female hoary squash bees were already exposed to doses above lethal levels of both imidacloprid and clothianidin (HQs >1) during the construction of a single nest, rendering it extremely unlikely they could construct another. Indeed, under present soil neonicotinoid residue conditions in Ontario *Cucurbita* cropping systems, pesticide exposure may preclude the construction of even a single 5 cell nest in a season even if all other conditions are favourable. Excluding exposure from nectar, which was not evaluated, hazard to larval stages was low (HQ_larvae,pollen_ = 0.06). It is perhaps reasonable to expect that hazard from nectar is also low for larval stages because their consumption of nectar is much less than for adult females as they are not involved with energy-expensive nest construction or foraging activities (HQ_nectar,adult female_ = 0.45). For honey bees, Rortais *et al.*^40^ reported that larvae consume 150% less nectar than adult pollen foragers. In our study we have assumed that larval hoary squash bees are protected from direct exposure to neonicotinoids in soil by the waterproof nature of the nest cell lining^15^, an assumption that requires further critical investigation.

The percentage translocation of insecticide residues from soil to bees is currently unknown. Our initial assumption here was translocation at 100%, but in recognition of the uncertainty existing around translocation rates we also present exceedances for four alternative scenarios with lower rates (Table S8). Using a probabilistic approach for the hoary squash bee in the acute exposure scenario (48 h, 2.23 g soil; Fig. 2, Table S6) at 100% translocation, the probability of exceedance of the mean honey bee LC_50_ was below 5% for imidacloprid and clothianidin. However, for the solitary bee surrogate LC_50_ endpoint residues of both imidacloprid (31.2%) and clothianidin (28.3%) in soil exceeded 5% (Fig. 2, Table S6). In the chronic exposure scenario, the amount of exposure to soil (33.5 g soil, 30 days) was much greater, and exceedance for imidacloprid and clothianidin was greater than 5% for all lethal endpoints (Fig. 3A,B, Table S6). It is likely that at least some of the clothianidin found in *Cucurbita* crop soil is from the application of thiamethoxam to seeds^29^. Hilton *et al*.^30^ found that ∼3-46 % of the residues recovered from soil sampled more than 60 days after thiamethoxam application were the metabolite clothianidin. Further work on the fate of thiamethoxam in *Cucurbita* crop fields is needed to determine whether seed-applied thiamethoxam poses a risk to hoary squash bees via its metabolite, clothianidin. Exceedance for chlorantraniliprole was not greater than 5% for any exposure endpoint in either exposure scenario, suggesting that it did not pose a risk in *Cucurbita*-crop soils in 2016 (Fig. 3C, Table S6).

There were at least three issues generating uncertainty during our assessment of potential risk to hoary squash bees posed by neonicotinoid residues in soil. The lack of information about insecticide toxicity for this, or indeed any other solitary ground-nesting, species is a large knowledge gap. As the hoary squash bee is similar in size to the honey bee it may be well represented by the available toxicity data for honey bees, especially for contact exposure which tends to vary less among species than measures of oral exposure^37^. Secondly, although soil-applied neonicotinoids are known to elicit negative effects on Lepidoptera that pupate in soil^41^, Carabid beetles that live in soil at all life stages^42^, and Hexapoda, Collembola, Thysanoptera, and Coleoptera adults^43^, we could find no information on the extent to which insecticide residues in soil can pass through the cuticle or into spiracles of bees that burrow in soil. We have assumed a worst-case scenario in which all the soil residues are translocated during exposure, but this is unlikely, although neonicotinoids have relatively low organic carbon-water partition coefficients (Koc; Table S4)^44^. However, even at 10% translocation exceedances were greater than 5% for all imidacloprid endpoints, and also for the lowest honey bee LC_50_ and solitary bee surrogate LC_50_ for clothianidin, for chronic exposure in *Cucurbita* crop soils,. Assuming a 10% translocation in a chronic exposure scenario in field crops, exceedances were greatrer than 5% for all endpoints for exposure to clothianidin (Table S8). Lastly, there is a need for lower residue detection limits for soil to align them with lethal dose concentrations for some compounds (Table S3).

Confidence intervals for the EEDs in *Cucurbita-*crop soil are large due to limited sample size and high LODs. Interestingly, even if we consider the extreme high values for confidence intervals (where exceedances would be lowest), exceedance remains above 5% for the solitary bee surrogate LC_50_ for clothianidin and imidacloprid respectively in both the acute and chronic scenarios (Figs. 2,3). Despite its limitations, this study is significant because it represents the first evaluation of risk from insecticide residues in soil for any ground-nesting bee species.

About 70% of the solitary bee species in eastern Canada nest in the ground^45^, many of which are associated with agriculture, including species in the genera *Agapostemon, Andrena, Anthophora, Colletes, Eucera, Halictus, Lasioglossum, Megachile* and *Melissodes*^46,47^. Because there is little information about exposure to soil for most ground-nesting bee species, this study used exposure of the hoary squash bee to soil during nest construction as a model for other ground-nesting bees. The estimate of soil exposure of 33.5g for a hoary squash bee female is comparable to that for some other ground-nesting bee species. For example, total exposure to soil for *Andrena prunorum* females has been estimated at 30.23 g^48^, and the amount of soil excavated by the female alkali bee (*Nomia melanderi*) is estimated to be 26.3 ± 5.7 g^49^. The difference between the low-end estimate of soil excavated by *N. melanderi* (20.6 g) and hoary squash bees (33.5 g) could represent as much as 38.5% of the latter species’ exposure, highlighting the potential variability in exposure via soil across species and the limitations of the hoary squash bee model. Ground-nesting bee species vary greatly in size^12^, and many are much smaller than hoary squash bees. For ground-nesting solitary bees, tunnel diameter is related to bee size because bees excavate tunnels and nest cells large enough for themselves^12^. Although other solitary ground-nesting bee species may be appreciably larger or smaller than squash bees, the ratio of their body size to the volume of soil they excavate when building a nest may be similar, providing a possible future basis upon which to compare exposure for different solitary bee species. Smaller bees may have lower exposure to neonicotinoids in soil because they construct narrower tunnels and nest cells to fit their smaller bodies^12^, therefore contacting lower soil volumes overall.

However smaller bee species may be more sensitive to insecticide exposure based on body size and other factors^14,32,50^. Bee body size is not necessarily correlated to the depth that vertical tunnels in nests are excavated in soil^51^. Solitary bees are physiologically limited in their reproductive capacity and generally build 1-8 brood cells per nest^12^. Until more information emerges, the hoary squash bee is the best model available to evaluate risk from exposure to pesticide residues in soil for ground-nesting bees in general and provides a starting point to understand risk from insecticides residues in soil for ground-nesting bees.

Neonicotinoid residues detected in Ontario’s agricultural soils reflect variation in usage for different crops. For *Cucurbita* crops, imidacloprid and clothianidin were the most commonly detected neonicotinoids, with a single detection of thiamethoxam at an unquantifiable concentration. For field crops, clothianidin and thiamethoxam were more commonly detected. Exposure to clothianidin residues for bees nesting in field crop soils appeared to be ubiquitous and chronic: 96.34% of soil samples taken before spring planting contained clothianidin applied in the previous season. The probabilities of exceedance were high for all the exposure endpoints for both the acute (Fig. 5) and chronic (Fig. 6) exposure to clothianidin. As clothianidin-treated seeds are planted in a new cropping cycle, releasing more residues into the soil, these exceedances will likely increase. Ground-nesting bees from 13 genera have been collected in corn and soybean fields in Iowa^52,53^. Although this does not prove that these species were nesting within fields, the small foraging ranges of solitary bees^54^ suggest that many likely were. Various bees are active at different times during the season^45,55^. Those species active in the early spring may be exposed to the minimum residue concentrations described here, but those active post-planting may be exposed to much higher concentrations in soil.

Taken together, this evidence suggests the overall risk to ground-nesting bees from exposure to clothianidin in field crop soil is high, necessitating action to mitigate such risks to preserve pollination services. If clothianidin residues found in soil originate as a metabolite of applied thiamethoxam, then use of thiamethoxam should also be addressed.

Thiamethoxam is used as a seed treatment on both field corn and soybean crops in Ontario and was applied to 1.98 million acres (66% of treated acres) in the 2016 season^56^. For the chronic exposure scenario, the risk to ground-nesting bees from thiamethoxam was high for all exposure endpoints (Fig. 5B). For the acute exposure scenario, the risk from thiamethoxam was less than 5% for all exposure endpoints except the solitary bee surrogate LC_50_ (25.56 %: Fig. 4). The apparently lower risk to ground-nesting bees from thiamethoxam may be the product of its tendency to break down quickly into clothianidin in soil^29^. Although the risk to ground-nesting bees from acute exposure to imidacloprid in field crop soil was below the 5% threshold for all exposure endpoints (Fig. S1), exceedance rose above the threshold for the solitary bee surrogate LC_50_ (9.24 %) under chronic exposure (Fig. S2). The lower risk associated with imidacloprid in field crop soil may be because it is used in only 11% of treated field crop acres^56^. One of the main concerns around neonicotinoid insecticide exposure for ground-nesting bees is their use in soil applications as treated seed. This has been partially mitigated in Ontario by increased regulation of neonicotinoid-treated corn and soybean seed^57^ but has not yet been addressed for other crops.

In conclusion, neonicotinoid residues in soil pose a high risk to female hoary squash bees as they construct their nests in *Cucurbita*-crop growing systems or in field crop soils. These demonstrable risks for hoary squash bees seem likely to be applicable to other species of ground-nesting bees nesting in agricultural soils. Further work is needed to determine the realative sensitivity of the hoary squash bee to neonicotinoid exposure compared to honey bees, and to explicitly determine the extent and impacts of larval exposure in soil. Advances in analytical techniques are also needed to achieve lower limits of detection in soil that mirror lethal endpoints for solitary bees. Recognition and mitigation of risks from exposure to neonicotinoids in agricultural soil are urgently needed to protect these important crop pollinators.

## Acknowledgements

We would like to thank all 18 Ontario farmers who allowed us access to their land and crops, Beatrice Chan and Katie Fisher for their assistance with field sampling, Linda Lissemore for advice on residue analyses, Elaine Roddy for providing input into production practices and pesticide use patterns on farms, providing farm contacts, and sampling on all the farms in southwestern Ontario, and Jim Chaput for technical advice on pesticide registration and use in *Cucurbita* crops. This work was supported by the Ontario Ministry of Agriculture, Food and Rural Affairs (OMAFRA) grant UofG2015-2466 (awarded to N.E.R and D.S.W.C.), the Ontario Ministry of Environment and Climate Change (MOECC) Best in Science grant BIS201617-06 (awarded to N.E.R.), Natural Sciences and Engineering Research Council (NSERC) Discovery grants 2015-06783 and 2018-04641 (awarded to N.E.R. and R.S.P. respectively), the Fresh Vegetable Growers of Ontario (FVGO: awarded to N.E.R and D.S.W.C.), and the Food from Thought: Agricultural Systems for a Healthy Planet Initiative, by the Canada First Research Excellent Fund (grant 000054). D.S.W.C. was supported by the George and Lois Whetham Scholarship in Food Systems, an Ontario Graduate Fellowship, the Keith and June Laver Scholarship in Horticulture, the Fred W. Presant Scholarship, and a Latournelle Travel Scholarship. N.E.R. is supported as the Rebanks Family Chair in Pollinator Conservation by The W. Garfield Weston Foundation.

## Author Contributions

D.S.W.C., R.S.P. and N.E.R. conceived and designed the project. D.S.W.C. carried out the experimental work and collated MOECC datasets. D.S.W.C., R.S.P. and J.L.R-G. carried out the probabilistic risk assessments and statistical analyses. All authors contributed to writing the paper.

