## Supplemental Information for "Assessment of risk to hoary squash bees (*Peponapis pruinosa*) and other ground-nesting bees from systemic insecticides in agricultural soil"

**Electonic Supplementary Information**

D. Susan Willis Chan^1^, Ryan S. Prosser^1^, Jose L. Rodríguez-Gil^2^, Nigel E. Raine^1^

^1^ School of Environmental Sciences, University of Guelph, Guelph, Ontario, N1G 2W1, Canada

^2^ Department of Biology, University of Ottawa, Ottawa, Ontario, K1N 6N5, Canada

*

**Figure S1**. Environmental Exposure Distribution (EED) for acute exposure to imidacloprid in soil for solitary ground-nesting bees based on MOECC data taken from field crop soils in southern Ontario, 2016. Soil samples taken from 0-15 cm depth. Effects benchmark concentrations (i.e. honey bee geometric mean LC_50_, solitary bee surrogate LC_50_) for acute exposure using the hoary squash bee (Peponapis pruinosa) model (30 days, 33.5 g soil) are represented by vertical lines on the EED. Exceedance of these endpoints is calculated by subtracting the cumulative probability from 1.0. Grey horizontal lines represent individual samples below the limit of detection.

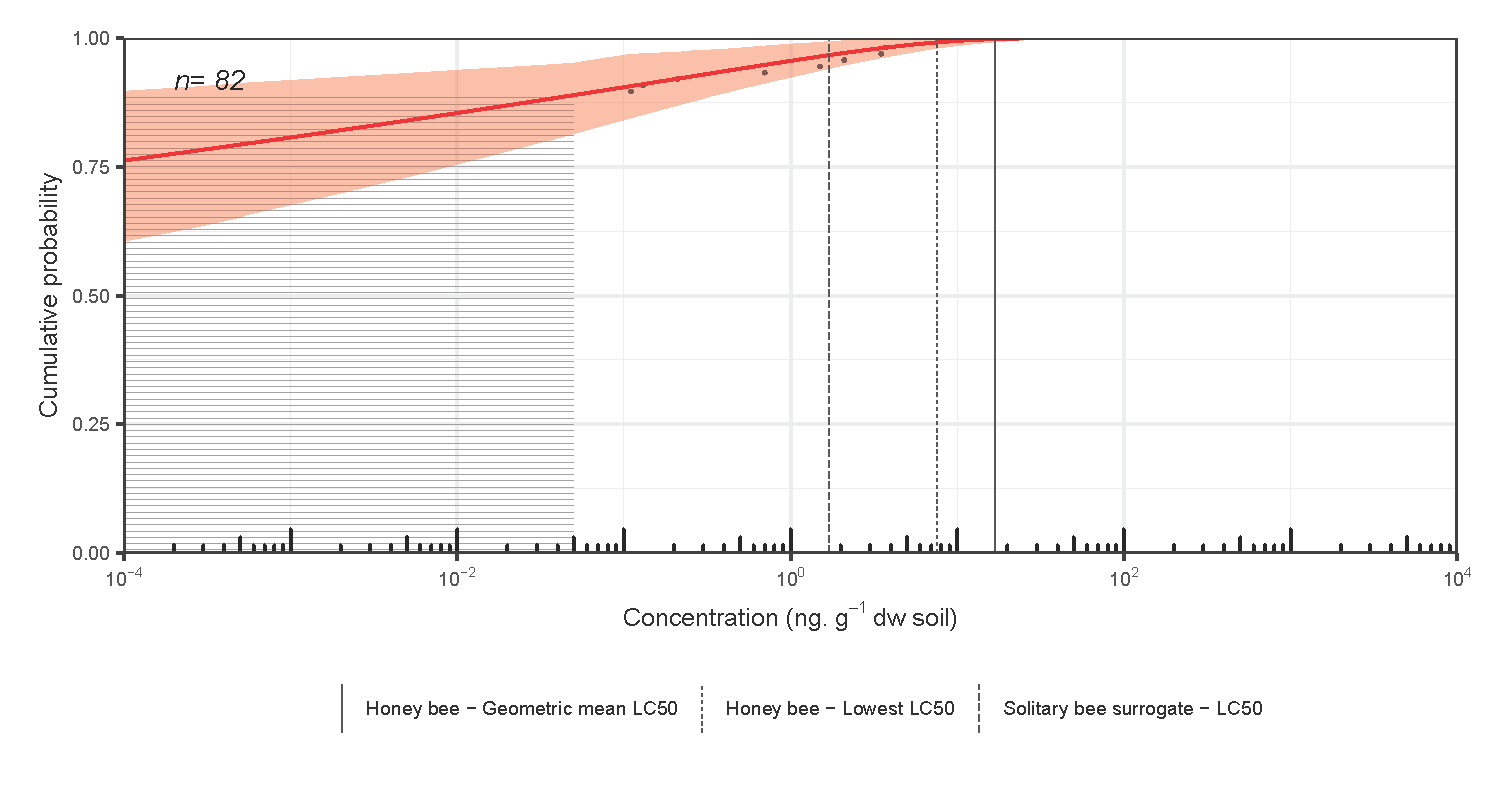

**Figure S2**. Environmental Exposure Distribution (EED) for chronic exposure to imidacloprid in soil for solitary ground-nesting bees based on MOECC data taken from field crop soils in southern Ontario, 2016. Soil samples taken from 0-15 cm depth. Effects benchmark concentrations (i.e. honey bee geometric mean LC_50_, solitary bee surrogate LC_50_) for chronic exposure using the hoary squash bee (Peponapis pruinosa) model (30 days, 33.5 g soil) are represented by vertical lines on the EED. Exceedance of these endpoints is calculated by subtracting the cumulative probability from 1.0. Grey horizontal lines represent individual samples below the limit of detection.

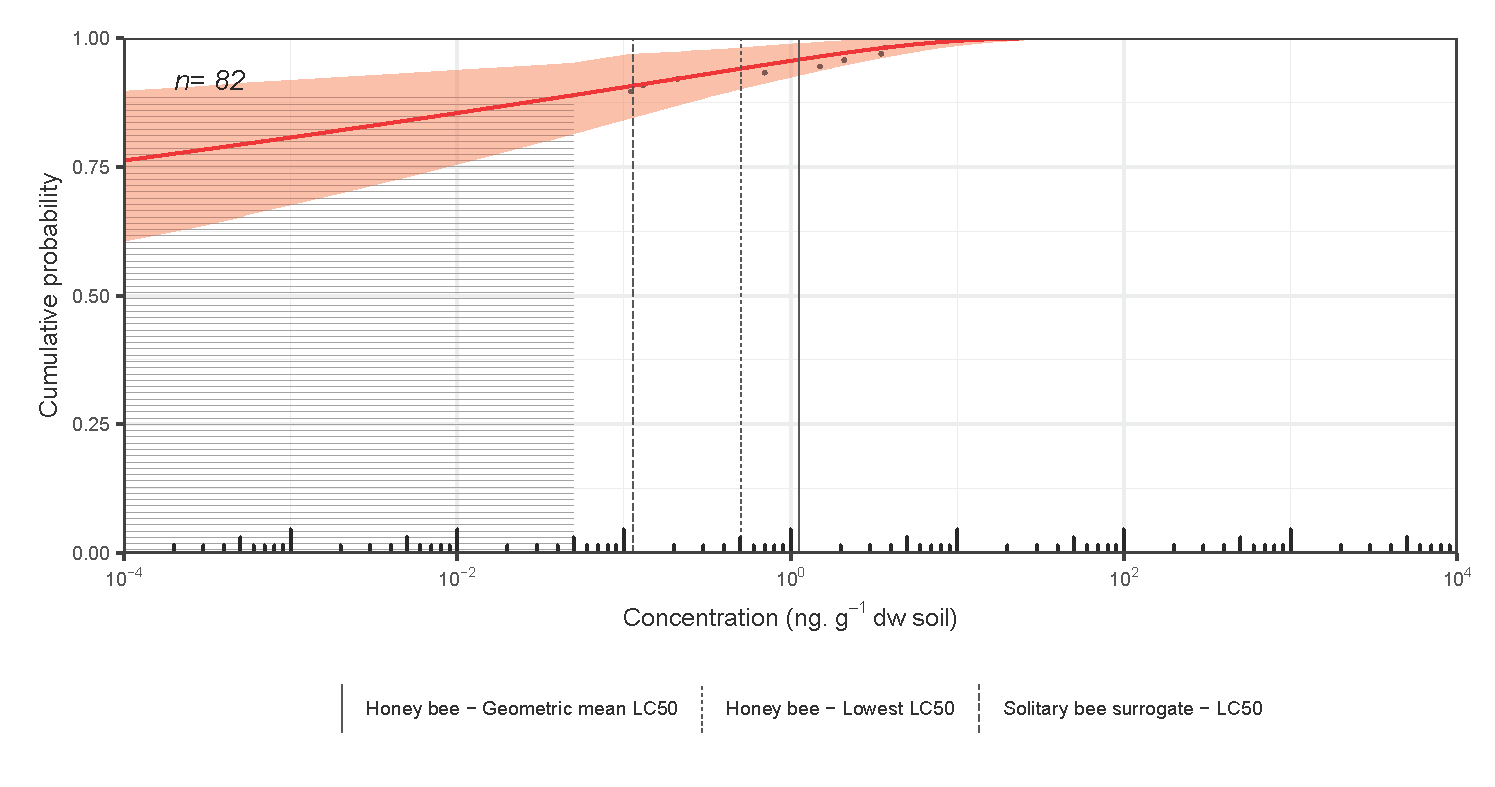

**Table S1**. Calculations of the volume, and therefore, mass of soil excavated by a female hoary squash bee (*Peponapis pruinosa*) to construct an underground nest, from which to determine potential insecticide exposure via soil. Nest dimensions follow data from Mathewson (1968)^1^.

| **Part of Nest** | **Soil volume/ mass** |
| --- | --- |
| a. Main vertical shaft  (7 mm diameter x 18 cm long) | 6.92 cm^3^ |
| b. Antechamber  (7 mm diameter x 6 cm long) | 2.31 cm^3^ |
| c. 5 brood cells  (7 mm diameter x 7.3 cm long = 2.81 cm^3^ each) | 14.04 cm^3^ |
| d. 5 brood caps  (7 mm diameter x 1 cm thick = 0.3847 cm^3^ each)  Estimated from our own observations | 1.92 cm^3^ |
| Total volume of soil excavated  (a + b + c + d) | 25.19 cm^3^ |
| Total mass of soil excavated  (Total volume x Bulk density (BD) of loam; BD = 1.33 g/cm^3^) | **33.51 g** |

**Table S2**. Summary of pesticide exposure routes, exposure types, exposure amount, and exposure period for the hoary squash bee (*Peponapis pruinosa*).

| **Exposure Route** | **Developmental Stage** | **Exposure type** | **Exposure amount** | **Period of Exposure** |
| --- | --- | --- | --- | --- |
| Soil | Adult Female | Contact | 33.51 g ^1^ | Construction of 1 nest  ~30 days |
| Soil | Larva | Contact | Putatively not exposed ^1^ | 10 months |
| Nectar | Adult Female | Oral | <<780 mg; based on pollen-foraging honey bee ^2^ | 30+ days |
| Nectar | Adult Male | Oral | Unknown, <adult female | 30+ days |
| Nectar | Larva | Oral | Unknown, < adult females | 15 days |
| Pollen | Larva | Oral | 54.2 mg ^3*^ | 15 days |
| Pollen | Larva | Contact | Unknown | 15 days |
| Pollen | Adult Female | Contact | 246 mg (5x larval exposure) | 30 days |
| Pollen | Adult Female | Oral | Unknown | During oocyte maturation^4^ |

* = present study

**Table S3**. All residues detected in soil, pollen, and nectar of *Cucurbita*-crop growing systems in Ontario, Canada, 2016, showing limits of detection and quantification (LOD/LOQ), maximum concentrations, geometric mean concentrations, and frequency of detection for each residue (n = 29 samples, nd = not detected, dnq=detected, not quantifiable). Frequency of detection is based on samples in which residues were detectable but not necessarily quantifiable (>LOD).

|  |  | **Soil** | | | | **Pollen** | | | | **Nectar** | | | |
| --- | --- | --- | --- | --- | --- | --- | --- | --- | --- | --- | --- | --- | --- |
|  |  | **LOD/ LOQ** | **Max Conc.** | **Mean Conc.** | **Freq. Detection** | **LOD/ LOQ** | **Max Conc.** | **Mean Conc.** | **Freq. Detection** | **LOD/**  **LOQ** | **Max Conc.** | **Mean Conc.** | **Freq. Detection** |
| **Type of pesticide** | **Active ingredient** | ppb | ng a.i./g matrix | ng a.i./g matrix | % | ppb | ng a.i./g matrix | ng a.i./g matrix | % | ppb | ng a.i./g matrix | ng a.i./g matrix | % |
| **Insecticide** | Clothianidin | 1/4 | 5.9 | 2.0 | 34 | 2/8 | nd | nd | 0 | 1/4 | nd | nd | 0 |
|  | Imidacloprid | 3/9 | 41.6 | 3.0 | 21 | 2/8 | 4.3 | 4.3 | 3 | 1/4 | 1.1 | 0.9 | 3 |
|  | Thiamethoxam | 5/20 | dnq | - | 3 | 2/8 | nd | nd | 0 | 0.6/2 | nd | nd | 0 |
|  | Chlorantraniliprole | 6/20 | 148.5 | 36.8 | 24 | 3/10 | 68 | 68 | 3 | 2/6 | nd | nd | 0 |
|  | Carbaryl | 1/4 | 352.8 | 14.2 | 10 | 2/7 | 31.1 | 16.5 | 7 | 0.6/2 | nd | nd | 0 |
|  | Methomyl | 7/20 | nd | - | 0 | 2/6 | nd | nd | 0 | 0.3/0.9 | 0.5 | 0.4 | 7 |
|  | Dimethoate | 2/6 | nd | - | 0 | 0.5/2 | 6.2 | 6.2 | 3 | 0.4/1 | 0.5 | 0.5 | 3 |
| **Fungicide** | Pyraclostrobin | 4/10 | 16.7 | 3.8 | 10 | 1/4 | 399.6 | 29.6 | 7 | 0.4/1 | 2 | 2 | 3 |
|  | Picoxystrobin | 0.4/1 | nd | nd | 0 | 0.7/2 | 110.2 | 4.6 | 10 | 0.2/0.7 | 0.3 | 0.3 | 3 |
|  | Boscalid | 3/9 | 374.9 | 46.2 | 31 | 2/6 | 25 | 17.8 | 7 | 2/8 | nd | nd | 0 |
|  | Propamocarb | 20/50 | 64.4 | 23.0 | 10 | 0.9/3 | 402.6 | 222.1 | 14 | 0.2/0.5 | 74.5 | 11.2 | 17 |
|  | Quinoxyfen | 5/10 | 14.7 | 7.9 | 3 | 3/8 | 94.9 | 79.1 | 7 | 1/3 | nd | nd | 0 |
|  | Difenoconazole | 4/10 | 40.7 | 18.9 | 14 | 2/5 | 22.4 | 16.5 | 7 | 0.7/2 | nd | nd | 0 |
| **Herbicide** | Napropamide | 1/4 | 59.4 | 2.8 | 7 | 0.4/1 | 5.0 | 5.0 | 3 | 0.1/0.4 | nd | nd | 0 |
|  | Linuron | 5/10 | 0.8 | 0.8 | 0 | 1/4 | nd | nd | 0 | 0.8/2 | nd | nd | 0 |

**Table S4**. Physical, chemical, and environmental fate properties of imidacloprid, clothianidin, thiamethoxam and chlorantraniliprole

| **Active Ingredient** | **Imidacloprid** | **Clothianidin** | **Thiamethoxam** | **Chlorantraniliprole** |
| --- | --- | --- | --- | --- |
| Chemical Name^a^ | 1-[(6-Chloro-3-pyridinyl)methyl]-N-nitro-2-imidazolidinimine | (E)-1-(2-chloro-1, 3-thiazol-5-ylmethyl)-3-methyl-2-nitroguanidine | 3-[(2-chloro-5-thiazolyl) methyl]tetrahydro-5-methyl-N-nitro-4H-1,3,5-oxadiazin-4-imine | 3-bromo-N-(4-chloro-2-methyl-6-((methylamino) carbonyl)phenyl)-1-(3-chloro-2-pyridinyl)-1H-pyrazole-5-carboxamide |
| Insecticide class | Neonicotinoid | Neonicotinoid | Neonicotinoid | Anthranilic diamide |
| Formula^a^ | C_9_H_10_ClN_5_O_2_ | C_6_N_5_H_8_SO_2_Cl | ‎C_8_H_10_ClN_5_O_3_S | C18-H14-Br-Cl2-N5-O2 |
| CAS No.^a^ | 138261-41-3 | 210880-92-5 | 153719-23-4 | 500008-45-7 |
| Molecular Weight^1^ (g/mol) ^a^ | 255.69 | 327 | 297.1 | 483.2 |
| Solubility in water at 20° C, pH 7 (mg/L) ^a^ | 610 ^5^ | 327 ^6^ | 4100 ^5^ | 0.9-1.0 ^5^ |
| Adsorption to Particles (soil) Koc at pH 7 ^a^ | 156-800 | 60 | 68.4 | 244-468 |
| Volatility (air)  Vapour pressure ^a^  (mm Hg at 25°C) | 7.0x10^-12^ | 9.8x10^-10^ | 4.95x10^-11^ | 1.2x10^-14^ |
| Log Kow ^a^ | 0.57 | 0.7 | -0.13 | 2.76 |
| Bio-degradation DT50 (days) | 48-190 ^5^ | 148-1155 ^5^ | 46.3-301 ^7^ | 60-365 |

**Table S5**. Exposure endpoints for amounts of soil handled by the hoary squash bee in acute (48h, 2.23 g soil) and chronic (30 days, 33.5 g soil) exposure scenarios based on various effect endpoints in the scientific literature^8-16^. HB = honey bee, SB = solitary bee.

| **Insecticide** | **Exposure type** | **Effect endpoint** | **Effect endpoint concentration (ng a.i./bee)**  **A** | **Exposure amount (g soil/bee)**  **B** | **Exposure endpoint**  **(A/B)**  **(ng a.i./g soil)** | **Source for effect endpoint** |
| --- | --- | --- | --- | --- | --- | --- |
| **Clothianidin** | Acute | Geomean HB LD_50_ | 35.88 | 2.23 | 15.16 | 8-11 |
|  |  | Lowest HB LD_50_ | 22 | 2.23 | 9.29 | 9 |
|  |  | SB Surrogate LD_50_ | 3.588 | 2.23 | 1.52 | 12 |
|  | Chronic | Geomean HB LD_50_ | 35.88 | 33.5 | 1.01 | 8-11 |
|  |  | Lowest HB LD_50_ | 22 | 33.5 | 0.62 | 9 |
|  |  | SB Surrogate LD_50_ | 3.588 | 33.5 | 0.10 | 12 |
| **Imidacloprid** | Acute | Geomean HB LD_50_ | 40.03 | 2.23 | 16.91 | 8,9,11,12,14 |
|  |  | Lowest HB LD_50_ | 18 | 2.23 | 7.61 | 9 |
|  |  | SB Surrogate LD_50_ | 4.003 | 2.23 | 1.69 | 12 |
|  | Chronic | Geomean HB LD_50_ | 40.03 | 33.5 | 1.13 | 8,9,11,13,14 |
|  |  | Lowest HB LD_50_ | 18 | 33.5 | 0.51 | 9 |
|  |  | SB Surrogate LD_50_ | 4.003 | 33.5 | 0.113 | 12 |
| **Thiamethoxam** | Acute | Geomean HB LD_50_ | 25.64 | 2.23 | 10.83 | 8,10,11,15 |
|  |  | Lowest HB LD_50_ | 24 | 2.23 | 10.14 | 8 |
|  |  | SB Surrogate LD_50_ | 2.564 | 2.23 | 1.08 | 12 |
|  | Chronic | Geomean HB LD_50_ | 25.64 | 33.5 | 0.722 | 8,10,11,15 |
|  |  | Lowest HB LD_50_ | 24 | 33.5 | 0.676 | 8 |
|  |  | SB Surrogate LD_50_ | 2.564 | 33.5 | 0.072 | 12 |
| **Chlorantraniliprole** | Acute | Lowest HB LD_50_ | >87,500 | 2.23 | 36547.09 | 16 |
|  |  | SB Surrogate LD_50_ | >8750 | 2.23 | 3654.71 | 12,16 |
|  | Chronic | Lowest HB LD_50_ | >87,500 | 33.5 | 2432.84 | 16 |
|  |  | SB Surrogate LD_50_ | >8750 | 33.5 | 243.28 | 12,16 |

**Table S6**. Exceedance probabilities (i.e. the frequency that effect endpoints were exceeded) for 100% translocation of residues from soil to bee, with upper and lower limits of the 95% confidence interval and the exposure concentrations associated with each effect endpoint, for all systemic insecticides detected in soil of *Cucurbita* and other agricultural field crops in Ontario for both chronic (30 days, 33.5 g soil) and acute (48h, 2.23 g soil) exposure scenarios. HB = honey bee, SB = solitary bee. Data for field crop soils from MOECC data^18^.

| **Insecticide** | **Exposure type** | **Crop system** | **Effect endpoint** | **% Exceedance** | **Lower limit of 95% CI** | **Upper limit of 95% CI** | **Effect concentration**  **ng ai/g soil** |
| --- | --- | --- | --- | --- | --- | --- | --- |
| Clothianidin | Chronic | Cucurbita | Lowest HB LC_50_ | 44.3 | 65.9 | 24.5 | 0.6 |
| Clothianidin | Chronic | Cucurbita | Geomean LC_50_ | 35.8 | 54.9 | 17.7 | 1.0 |
| Clothianidin | Chronic | Cucurbita | SB Surrogate LD_50_ | 68.7 | 89.1 | 41.5 | 0.1 |
| Clothianidin | Acute | Cucurbita | Lowest HB LC_50_ | 2.4 | 4.6 | 0.4 | 9.3 |
| Clothianidin | Acute | Cucurbita | Geomean LC_50_ | 0.5 | 1.4 | 0.04 | 15.2 |
| Clothianidin | Acute | Cucurbita | SB Surrogate LD_50_ | 28.3 | 44.5 | 14.2 | 1.5 |
| Imidacloprid | Chronic | Cucurbita | Lowest HB LC_50_ | 57.8 | 100 | 26.2 | 0.5 |
| Imidacloprid | Chronic | Cucurbita | Geomean LC_50_ | 39.8 | 99.9 | 17.2 | 1.1 |
| Imidacloprid | Chronic | Cucurbita | SB Surrogate LD_50_ | 85.4 | 100 | 45.4 | 0.1 |
| Imidacloprid | Acute | Cucurbita | Lowest HB LC_50_ | 8.9 | 17.0 | 0.0 | 7.6 |
| Imidacloprid | Acute | Cucurbita | Geomean LC_50_ | 3.5 | 9.0 | 0.0 | 16.9 |
| Imidacloprid | Acute | Cucurbita | SB Surrogate LD_50_ | 31.2 | 95.6 | 13.7 | 1.7 |
| Chlorantraniliprole | Chronic | Cucurbita | Lowest HB LC_50_ | 0.0 | 0.0 | 0.0 | 2432.8 |
| Chlorantraniliprole | Chronic | Cucurbita | SB Surrogate LD_50_ | 1.6 | 3.4 | 0.12 | 243.3 |
| Chlorantraniliprole | Acute | Cucurbita | Lowest HB LC_50_ | 0.0 | 0 | 0 | 36547.1 |
| Chlorantraniliprole | Acute | Cucurbita | SB Surrogate LD_50_ | 0.0 | 0.0 | 0 | 3654.7 |
| Clothianidin | Chronic | Field crops | Lowest HB LC_50_ | 92.4 | 97.0 | 86.0 | 0.6 |
| Clothianidin | Chronic | Field crops | Geomean LC_50_ | 87.7 | 94.0 | 80.1 | 1.0 |
| Clothianidin | Chronic | Field crops | SB Surrogate LD_50_ | 98.8 | 99.8 | 96.5 | 0.1 |
| Clothianidin | Acute | Field crops | Lowest HB LC_50_ | 27.3 | 35.3 | 19.1 | 9.3 |
| Clothianidin | Acute | Field crops | Geomean LC_50_ | 11.7 | 17.6 | 5.8 | 15.2 |
| Clothianidin | Acute | Field crops | SB Surrogate LD_50_ | 81.9 | 89.7 | 73.6 | 1.5 |
| Imidacloprid | Chronic | Field crops | Lowest HB LC_50_ | 5.9 | 9.7 | 2.1 | 0.5 |
| Imidacloprid | Chronic | Field crops | Geomean LC_50_ | 4.2 | 7.1 | 1.3 | 1.1 |
| Imidacloprid | Chronic | Field crops | SB Surrogate LD_50_ | 9.2 | 15.3 | 3.9 | 0.1 |
| **Insecticide** | **Exposure type** | **Crop system** | **Effect endpoint** | **% Exceedance** | **Lower limit of 95% CI** | **Upper limit of 95% CI** | **Effect concentration**  **ng ai/g soil** |
| Imidacloprid | Acute | Field crops | Lowest HB LC_50_ | 0.8 | 1.8 | 0.0 | 7.6 |
| Imidacloprid | Acute | Field crops | Geomean LC_50_ | 0.2 | 0.7 | 0.0 | 16.9 |
| Imidacloprid | Acute | Field crops | SB Surrogate LD_50_ | 3.3 | 5.6 | 0.9 | 1.7 |
| Thiamethoxam | Chronic | Field crops | Lowest HB LC_50_ | 37.4 | 45.8 | 29.4 | 0.7 |
| Thiamethoxam | Chronic | Field crops | Geomean LC_50_ | 35.7 | 44.0 | 27.8 | 0.7 |
| Thiamethoxam | Chronic | Field crops | SB Surrogate LD_50_ | 78.4 | 85.8 | 69.7 | 0.1 |
| Thiamethoxam | Acute | Field crops | Lowest HB LC_50_ | 0.0 | 0.3 | 0.0 | 10.1 |
| Thiamethoxam | Acute | Field crops | Geomean LC_50_ | 0.0 | 0.2 | 0.0 | 10.8 |
| Thiamethoxam | Acute | Field crops | SB Surrogate LD_50_ | 25.6 | 33.1 | 17.8 | 1.1 |

**Table S7.** Parameters (shape and rate) and associated 95% confidence interval for the gamma (clothianidin, chlorantraniliprole in *Cucurbita* crops; all residues in field crops) or log-normal (imidacloprid, thiamethoxam in Cucurbita crops) model that was fit to the distribution of measured concentrations of each insecticide and the Akaike Information Criterion (AIC) for each model.

| **Insecticide** | **Sample depth/ cm** | **Crop type** | **shape** | **shape_2.5** | **shape_97.5** | **rate** | **rate_2.5** | **rate_97.5** | **AIC** |
| --- | --- | --- | --- | --- | --- | --- | --- | --- | --- |
| Clothianidin | 0-15 | Cucurbita | 0.34304 | 0.14723 | 0.66789 | 0.22847 | 0.13871 | 0.39616 | 76.905 |
| Imidacloprid | 0-15 | Cucurbita | -0.05037 | -2.51720 | 0.91093 | 1.58923 | 0.21634 | 3.15571 | 56.837 |
| Chlorantraniliprole | 0-15 | Cucurbita | 0.09636 | 0.03414 | 0.20079 | 0.00521 | 0.00285 | 0.01329 | 107.633 |
| Clothianidin | 0-15 | Field crops | 1.06754 | 0.75564 | 1.53303 | 0.15031 | 0.09975 | 0.23856 | 508.432 |
| Imidacloprid | 0-15 | Field crops | 0.02543 | 0.01082 | 0.04471 | 0.10976 | 0.04719 | 0.39991 | 91.718 |
| Thiamethoxam | 0-15 | Field crops | 0.53550 | 0.39736 | 0.73792 | 0.64797 | 0.43633 | 1.09491 | 205.267 |

**Table S8.** Exceedance of 3 effect endpoints (honey bee geometric mean LD_50_, honey bee lowest LD_50_, solitary bee surrogate LD_50_) for less than 100% translocation (75%, 50%, 25%, 10%) of neonicotinoid insecticide (clothianidin, imidacloprid, thiamethoxam) residue from soil to female ground-nesting bees as they construct nests in either an acute (48 h, 2.23 g soil) or chronic (30 days, 33.5 g soil) exposure scenario for two crop types (*Cucurbita* crops, field crops) based on exposure for the hoary squash bee (*Peponapis pruinosa*).

| **Crop type** | **Compound** | **Exposure Scenario** | **Effect Endpoint** | | **Trans-location**  **(%)** | | **Exceedance (%)** | | **Lower 95% CI** | | **Upper 95% CI** | | **Effect Concentration (ng ai/g soil)** |
| --- | --- | --- | --- | --- | --- | --- | --- | --- | --- | --- | --- | --- | --- |
| Cucurbita | Clothianidin | Acute | Honey bee Geometric mean LC_50_ | | 10% | | 0 | | 0 | | 0 | | 151.6052 |
|  |  |  |  | | 25% | | 0 | | 0 | | 0 | | 60.6421 |
|  |  |  |  | | 50% | | 0.01 | | 0.1 | | 0 | | 30.3210 |
|  |  |  |  | | 75% | | 0.15 | | 0.52 | | 0.01 | | 20.2140 |
|  |  |  | Honey bee Lowest LC_50_ | | 10% | | 0 | | 0 | | 0 | | 92.9577 |
|  |  |  |  | | 25% | | 0 | | 0.03 | | 0 | | 37.1831 |
|  |  |  |  | | 50% | | 0.22 | | 0.71 | | 0.01 | | 18.5915 |
|  |  |  |  | | 75% | | 1.07 | | 2.39 | | 0.12 | | 12.3944 |
|  |  |  | Solitary bee Surrogate LC_50_ | | 10% | | 0.53 | | 1.35 | | 0.04 | | 15.1607 |
|  |  |  |  | | 25% | | 6.08 | | 10.14 | | 1.80 | | 6.0643 |
|  |  |  |  | | 50% | | 15.88 | | 25.77 | | 7.21 | | 3.0321 |
|  |  |  |  | | 75% | | 23.01 | | 36.46 | | 11.10 | | 2.0214 |
|  |  | Chronic | Honey bee Geometric mean LC_50_ | | 10% | | 1.96 | | 3.85 | | 0.31 | | 10.1070 |
|  |  |  |  | | 25% | | 11.33 | | 18.8 | | 4.45 | | 4.0428 |
|  |  |  |  | | 50% | | 23.01 | | 36.46 | | 11.1 | | 2.0214 |
|  |  |  |  | | 75% | | 30.48 | | 47.55 | | 15.21 | | 1.3476 |
|  |  |  | Honey bee Lowest LC_50_ | | 10% | | 5.84 | | 9.75 | | 1.72 | | 6.1972 |
|  |  |  |  | | 25% | | 19.35 | | 31.18 | | 9.19 | | 2.4789 |
|  |  |  |  | | 50% | | 32.03 | | 49.68 | | 15.92 | | 1.2394 |
|  |  |  |  | | 75% | | 39.37 | | 59.67 | | 19.62 | | 0.8263 |
| Table S8 Continued | | | | | | | | | | | | | |
| **Crop type** | **Compound** | **Exposure Scenario** | **Effect Endpoint** | | **Trans-location**  **(%)** | | **Exceedance (%)** | | **Lower 95% CI** | | **Upper 95% CI** | | **Effect Concentration (ng ai/g soil)** |
| Cucurbita | Clothianidin | Chronic | Solitary bee Surrogate LC_50_ | | 10% | | 35.78 | | 54.9 | | 17.68 | | 1.0107 |
|  |  |  |  | | 25% | | 51.13 | | 73.42 | | 29.67 | | 0.4043 |
|  |  |  |  | | 50% | | 60.79 | | 82.91 | | 35.58 | | 0.2021 |
|  |  |  |  | | 75% | | 65.60 | | 86.87 | | 39.11 | | 0.1348 |
| Cucurbita | Imidacloprid | Acute | Honey bee Geometric mean LC_50_ | 10% | | 0.09 | | 1.19 | | 0 | | 169.1370 | |
|  |  |  |  | 25% | | 0.47 | | 2.80 | | 0 | | 67.6548 | |
|  |  |  |  | 50% | | 1.38 | | 4.99 | | 0 | | 33.8274 | |
|  |  |  |  | 75% | | 2.43 | | 7.00 | | 0 | | 22.5516 | |
|  |  |  | Honey bee Lowest LC_50_ | 10% | | 0.38 | | 2.51 | | 0 | | 76.0563 | |
|  |  |  |  | 25% | | 1.61 | | 5.50 | | 0 | | 30.4225 | |
|  |  |  |  | 50% | | 4.04 | | 9.82 | | 0 | | 15.2113 | |
|  |  |  |  | 75% | | 6.49 | | 13.38 | | 0 | | 10.1408 | |
|  |  |  | Solitary bee Surrogate LC_50_ | 10% | | 3.54 | | 8.96 | | 0 | | 16.9137 | |
|  |  |  |  | 25% | | 9.96 | | 18.47 | | 0 | | 6.7655 | |
|  |  |  |  | 50% | | 18.74 | | 32.09 | | 3.37 | | 3.3827 | |
|  |  |  |  | 75% | | 25.59 | | 66.38 | | 12.06 | | 2.2552 | |
|  |  | Chronic | Honey bee Geometric mean LC_50_ | 10% | | 5.76 | | 12.43 | | 0 | | 11.2758 | |
|  |  |  |  | 25% | | 14.64 | | 25.79 | | 0.18 | | 4.5103 | |
|  |  |  |  | 50% | | 25.59 | | 66.38 | | 12.06 | | 2.2552 | |
|  |  |  |  | 75% | | 33.57 | | 98.77 | | 14.41 | | 1.5034 | |
|  |  |  | Honey bee Lowest LC_50_ | 10% | | 13.17 | | 23.72 | | 0.03 | | 5.0704 | |
|  |  |  |  | 25% | | 27.58 | | 81.16 | | 12.64 | | 2.02817 | |
|  |  |  |  | 50% | | 42.10 | | 100 | | 18.75 | | 1.0141 | |
|  |  |  |  | 75% | | 51.29 | | 100 | | 23.08 | | 0.6761 | |
| Table S8 Continued | | | | | | | | | | | | | |
| **Crop type** | **Compound** | **Exposure Scenario** | **Effect Endpoint** | **Trans-location**  **(%)** | | **Exceedance (%)** | | **Lower 95% CI** | | **Upper 95% CI** | | **Effect Concentration (ng ai/g soil)** | |
| Cucurbita | Imidacloprid | Chronic | Solitary bee Surrogate LC_50_ | 10% | | 39.76 | | 99.98 | | 17.24 | | 1.1276 | |
|  |  |  |  | 25% | | 60.38 | | 100 | | 27.55 | | 0.4510 | |
|  |  |  |  | 50% | | 74.52 | | 100 | | 36.09 | | 0.2255 | |
|  |  |  |  | 75% | | 81.36 | | 100 | | 41.47 | | 0.1503 | |
| Field crops | Clothianidin | Acute | Honey bee Geometric mean LC_50_ | | 10% | | 0 | | 0 | | 0 | | 151.6052 |
|  |  |  |  | | 25% | | 0.02 | | 0.14 | | 0 | | 60.6421 |
|  |  |  |  | | 50% | | 1.31 | | 3.33 | | 0.21 | | 30.3210 |
|  |  |  |  | | 75% | | 5.65 | | 10.03 | | 1.91 | | 20.2140 |
|  |  |  | Honey bee Lowest LC_50_ | | 10% | | 0 | | 0 | | 0 | | 92.9577 |
|  |  |  |  | | 25% | | 0.48 | | 1.62 | | 0.04 | | 37.1831 |
|  |  |  |  | | 50% | | 7.14 | | 12.04 | | 2.74 | | 18.5915 |
|  |  |  |  | | 75% | | 17.47 | | 24.31 | | 10.26 | | 12.3944 |
|  |  |  | Solitary bee Surrogate LC_50_ | | 10% | | 11.72 | | 17.73 | | 5.64 | | 15.1607 |
|  |  |  |  | | 25% | | 43.23 | | 51.81 | | 34.93 | | 6.0643 |
|  |  |  |  | | 50% | | 66.33 | | 75.23 | | 57.83 | | 3.0321 |
|  |  |  |  | | 75% | | 76.35 | | 84.49 | | 67.76 | | 2.0214 |
|  |  | Chronic | Honey bee Geometric mean LC_50_ | | 10% | | 24.26 | | 31.72 | | 16.49 | | 10.1070 |
|  |  |  |  | | 25% | | 57.56 | | 66.27 | | 48.77 | | 4.0428 |
|  |  |  |  | | 50% | | 76.35 | | 84.49 | | 67.76 | | 2.0214 |
|  |  |  |  | | 75% | | 83.76 | | 90.73 | | 75.71 | | 1.3476 |
|  |  |  | Honey bee Lowest LC_50_ | | 10% | | 42.42 | | 50.97 | | 34.05 | | 6.1972 |
|  |  |  |  | | 25% | | 71.65 | | 80.37 | | 63.01 | | 2.4789 |
|  | Clothianidin | Chronic | Honey bee Lowest LC_50_ | | 50% | | 85.00 | | 91.72 | | 77.12 | | 1.2394 |
|  |  |  |  | | 75% | | 89.89 | | 95.21 | | 82.99 | | 0.8263 |
| Table S8 Continued | | | | | | | | | | | | | |
| **Crop type** | **Compound** | **Exposure Scenario** | **Effect Endpoint** | | **Trans-location**  **(%)** | | **Exceedance (%)** | | **Lower 95% CI** | | **Upper 95% CI** | | **Effect Concentration (ng ai/g soil)** |
| Field crops | Clothianidin | Chronic | Solitary bee Surrogate LC_50_ | | 10% | | 87.68 | | 93.67 | | 80.2 | | 1.0107 |
|  |  |  |  | | 25% | | 95.06 | | 98.30 | | 90.03 | | 0.4043 |
|  |  |  |  | | 50% | | 97.57 | | 99.40 | | 93.98 | | 0.2021 |
|  |  |  |  | | 75% | | 98.40 | | 99.68 | | 95.54 | | 0.1348 |
| Field crops | Imidacloprid | Acute | Honey bee Geometric mean LC_50_ | | 10% | | 0 | | 0 | | 0 | | 169.1370 |
|  |  |  |  | | 25% | | 0 | | 0 | | 0 | | 67.6548 |
|  |  |  |  | | 50% | | 0.02 | | 0.19 | | 0 | | 33.8274 |
|  |  |  |  | | 75% | | 0.08 | | 0.43 | | 0 | | 22.5516 |
|  |  |  | Honey bee Lowest LC_50_ | | 10% | | 0 | | 0 | | 0 | | 76.0563 |
|  |  |  |  | | 25% | | 0.03 | | 0.24 | | 0 | | 30.4225 |
|  |  |  |  | | 50% | | 0.23 | | 0.81 | | 0 | | 15.2113 |
|  |  |  |  | | 75% | | 0.51 | | 1.39 | | 0 | | 10.1408 |
|  |  |  | Solitary bee Surrogate LC_50_ | | 10% | | 0.18 | | 0.70 | | 0 | | 16.9137 |
|  |  |  |  | | 25% | | 0.94 | | 2.14 | | 0.03 | | 6.7655 |
|  |  |  |  | | 50% | | 1.97 | | 3.85 | | 0.24 | | 3.3827 |
|  |  |  |  | | 75% | | 2.75 | | 5.01 | | 0.47 | | 2.2552 |
|  |  | Chronic | Honey bee Geometric mean LC_50_ | | 10% | | 0.43 | | 1.24 | | 0 | | 11.2758 |
|  |  |  |  | | 25% | | 1.53 | | 3.16 | | 0.13 | | 4.5103 |
|  |  |  |  | | 50% | | 2.75 | | 5.01 | | 0.47 | | 2.2552 |
|  |  |  |  | | 75% | | 3.56 | | 6.25 | | 0.84 | | 1.5034 |
|  |  |  | Honey bee Lowest LC_50_ | | 10% | | 1.33 | | 2.83 | | 0.09 | | 5.0704 |
|  |  |  |  | | 25% | | 2.94 | | 5.28 | | 0.54 | | 2.0282 |
|  |  |  |  | | 50% | | 4.37 | | 7.50 | | 1.16 | | 1.0141 |
|  |  |  |  | | 75% | | 5.25 | | 8.81 | | 1.58 | | 0.6761 |
| Table S8 Continued | | | | | | | | | | | | | |
| **Crop type** | **Compound** | **Exposure Scenario** | **Effect Endpoint** | | **Trans-location**  **(%)** | | **Exceedance (%)** | | **Lower 95% CI** | | **Upper 95% CI** | | **Effect Concentration (ng ai/g soil)** |
| Field crops | Imidacloprid | Chronic | Solitary bee Surrogate LC_50_ | | 10% | | 4.16 | | 7.19 | | 1.07 | | 1.1276 |
|  |  |  |  | | 25% | | 6.14 | | 10.16 | | 2.00 | | 0.4510 |
|  |  |  |  | | 50% | | 7.69 | | 12.70 | | 2.65 | | 0.2255 |
|  |  |  |  | | 75% | | 8.60 | | 14.26 | | 2.95 | | 0.1503 |
| Field crops | Thiamethoxam | Acute | Honey bee Geometric mean LC_50_ | | 10% | | 0 | | 0 | | 0 | | 108.3263 |
|  |  |  |  | | 25% | | 0 | | 0 | | 0 | | 43.3305 |
|  |  |  |  | | 50% | | 0 | | 0 | | 0 | | 21.6653 |
|  |  |  |  | | 75% | | 0 | | 0.04 | | 0 | | 14.4435 |
|  |  |  | Honey bee Lowest LC_50_ | | 10% | | 0 | | 0 | | 0 | | 101.4084 |
|  |  |  |  | | 25% | | 0 | | 0 | | 0 | | 40.5634 |
|  |  |  |  | | 50% | | 0 | | 0 | | 0 | | 20.2817 |
|  |  |  |  | | 75% | | 0 | | 0.05 | | 0 | | 13.5211 |
|  |  |  | Solitary bee Surrogate LC_50_ | | 10% | | 0.02 | | 0.18 | | 0 | | 10.8326 |
|  |  |  |  | | 25% | | 2.02 | | 4.65 | | 0.4 | | 4.3331 |
|  |  |  |  | | 50% | | 10.37 | | 15.91 | | 4.87 | | 2.1665 |
|  |  |  |  | | 75% | | 18.71 | | 25.68 | | 11.15 | | 1.4444 |
|  |  | Chronic | Honey bee Geometric mean LC_50_ | | 10% | | 0.26 | | 1.07 | | 0.02 | | 7.2218 |
|  |  |  |  | | 25% | | 5.93 | | 10.26 | | 2.12 | | 2.8887 |
|  |  |  |  | | 50% | | 18.71 | | 25.68 | | 11.15 | | 1.4444 |
|  |  |  |  | | 75% | | 28.48 | | 36.27 | | 20.02 | | 0.9629 |
|  |  |  | Honey bee Lowest LC_50_ | | 10% | | 0.36 | | 1.37 | | 0.03 | | 6.7606 |
|  |  |  |  | | 25% | | 6.83 | | 11.45 | | 2.63 | | 2.7042 |
|  |  |  | Honey bee Lowest LC_50_ | | 50% | | 20.20 | | 27.31 | | 12.46 | | 1.3521 |
|  |  |  |  | | 75% | | 30.14 | | 38.01 | | 21.58 | | 0.9014 |
| Table S8 Continued | | | | | | | | | | | | | |
| **Crop type** | **Compound** | **Exposure Scenario** | **Effect Endpoint** | | **Trans-location**  **(%)** | | **Exceedance (%)** | | **Lower 95% CI** | | **Upper 95% CI** | | **Effect Concentration (ng ai/g soil)** |
| Field crops | Thiamethoxam | Chronic | Solitary bee Surrogate LC50 | | 10% | | 35.70 | | 43.76 | | 27.11 | | 0.7222 |
|  |  |  |  | | 25% | | 56.88 | | 65.8 | | 48.07 | | 0.2889 |
|  |  |  |  | | 50% | | 69.26 | | 77.86 | | 60.10 | | 0.1444 |
|  |  |  |  | | 75% | | 74.98 | | 83.11 | | 66.36 | | 0.0963 |
